# Telophase correction refines division orientation in stratified epithelia

**DOI:** 10.1101/668244

**Authors:** Kendall J. Lough, Kevin M. Byrd, Carlos P. Descovich, Danielle C. Spitzer, Abby J. Bergman, Gerard M. Beaudoin, Louis F. Reichardt, Scott E. Williams

## Abstract

During organogenesis, precise control of spindle orientation ensures a proper balance of proliferation and differentiation. In the developing murine epidermis, planar and perpendicular divisions yield symmetric and asymmetric fate outcomes, respectively. Classically, division axis specification involves centrosome migration and spindle rotation, events that occur early in mitosis. Here, we identify a previously uncharacterized orientation mechanism that occurs during telophase, correcting erroneous oblique orientations that unexpectedly persist into anaphase. The directionality of reorientation—towards either planar or perpendicular—correlates with the maintenance or loss of basal contact by the apical daughter. While the conserved scaffolding protein Pins/LGN is believed to function primarily through initial spindle positioning, we now show it also functions actively during telophase to reorient oblique divisions toward perpendicular. The ability to undergo telophase correction is also critically dependent upon an LGN-independent pathway involving the tension-sensitive adherens junction proteins vinculin, a-catenin and afadin, and correction directionality is influenced by local cell density. Failure of this reorientation mechanism impacts tissue architecture, as excessive oblique divisions induce precocious differentiation. The division orientation plasticity provided by telophase correction may provide a means for progenitors to dynamically respond to extrinsic cues provided by neighboring cells in order to adapt to local tissue needs.

## INTRODUCTION

Stem and progenitor cells utilize asymmetric cell divisions to balance self-renewal and differentiation. Cell fate decisions can be influenced by the division axis, with the choice between symmetric and asymmetric fate outcomes dictated by positioning of the mitotic spindle. Mechanistically, precise control of division orientation may serve to equally or unequally partition fate determinants, or restrict access to a stem cell niche (Knoblich, 2008; Siller & Doe, 2009). Errors in division orientation can lead to defects in differentiation and cell identity with the potential to drive overgrowths associated with cancer (Knoblich, 2010; Martin-Belmonte & Perez-Moreno, 2011; Neumuller & Knoblich, 2009).

The developing murine epidermis serves as an excellent model for studying how oriented cell divisions (OCDs) direct cell fate choices. Basal progenitors are capable of dividing either within the plane of the epithelium or perpendicular to it, resulting in symmetric or asymmetric divisions, respectively (Lechler & Fuchs, 2005; Smart, 1970). This process is governed by a conserved complex of spindle orienting proteins, including the essential linker LGN/Gpsm2 (Williams, Beronja, Pasolli, & Fuchs, 2011; Williams, Ratliff, Postiglione, Knoblich, & Fuchs, 2014). During epidermal and oral epithelial stratification, LGN is recruited to the apical cortex in ~50% of mitoses, and LGN loss leads to increased planar divisions and severe differentiation defects (Byrd et al., 2016; Williams et al., 2011; Williams et al., 2014). Thus, a parsimonious explanation for the observed bimodal distribution of division angles is that perpendicular divisions occur when sufficient levels of LGN are recruited to the apical cortex during early mitosis, and planar divisions occur when this apical recruitment fails.

In this and other models, it is assumed that the division axis is established relatively early in mitosis, either through directed centrosome migration or spindle rotation. As an example of the former, in the Drosophila melanogaster testis and larval neuroblasts, one centrosome migrates to the opposite side of the cell during prophase, and the metaphase spindle forms along, and remains fixed by, this centrosomal axis (Rebollo, Roldan, & Gonzalez, 2009; Siller, Cabernard, & Doe, 2006; Yamashita, Jones, & Fuller, 2003). In other systems—including the C. elegans early embryo, D. melanogaster embryonic neuroblasts, and progenitors of the vertebrate neuroepithelia—the spindle dynamically rotates during metaphase to align with extrinsic niche-derived or intrinsic polarity cues (Geldmacher-Voss, Reugels, Pauls, & Campos-Ortega, 2003; Haydar, Ang, & Rakic, 2003; Hyman & White, 1987; Kaltschmidt, Davidson, Brown, & Brand, 2000). Live imaging of cultured keratinocytes suggests that spindle orientation is fixed at metaphase by centrosome migration (Poulson & Lechler, 2010), but it is unclear whether this also holds true *in vivo*.

Here, utilizing *ex vivo* live imaging in combination with mosaic RNAi, we find that division orientation in the developing murine epidermis is not determined solely by LGN localization during prophase. In contrast, spindle orientation remains dynamic even into late stages of mitosis, and surprisingly, division axes remain random and uncommitted long after metaphase. While ~half of cells enter anaphase with planar (0-20°) or perpendicular (70-90°) orientations and maintain this division axis through telophase, the other half are initially oriented obliquely (20-70°), but undergo dramatic reorientation, a process we term telophase correction. In addition, we demonstrate that the α-catenin/vinculin/afadin cytoskeletal scaffolding complex is required for this correction to occur, and likely functions to modulate the tensile properties of the cell cortex by altering how actin is recruited to adherens junctions (AJs). These studies support a novel two-step model of OCD, where intrinsic factors such as LGN provide spatial cues that guide initial spindle positioning during early mitosis, while extrinsic factors such as cell-cell adhesions may provide a tension or density-sensing mechanism that provides flexibility to allow the division plane to reorient during telophase to adapt to tissue requirements. Our data further suggest that these mechanisms are modulated over developmental time to coordinate progenitor-expansive and differentiative programs.

## RESULTS

### Oblique anaphase divisions reorient during telophase

At embryonic day (E)16.5, during peak stratification, epidermal basal cells undergo either LGN-dependent perpendicular divisions or LGN-independent planar divisions, with roughly equal frequency. In support of this, LGN is invariantly apical when recruited to the cell cortex during prophase and remains apical at telophase in perpendicular divisions (Williams et al., 2011; Williams et al., 2014). However, both cortical LGN and the spindle axis can be observed at intermediate angles at metaphase (Williams et al., 2011), raising the question of when and how the spindle axis becomes fixed during epidermal OCDs.

As a first step to examine the dynamics of spindle positioning during peak stratification (E16.5), we characterized division orientation patterns at distinct stages of mitosis in fixed sections. To identify rare anaphase cells, we relied on the cleavage furrow marker survivin, which displays distinct localization patterns at anaphase and telophase, appearing disperse during the former and concentrating into two puncta during the latter (Williams et al., 2011) (Fig. 1A). In agreement with our previous observation, metaphase spindles were oriented randomly, suggesting that some degree of rotation occurs during metaphase. Surprisingly, however, the distribution of division angles remained random at anaphase, only establishing a bimodal distribution in telophase (Fig. 1B,C). These data suggest that active spindle orientation mechanisms occur after anaphase onset.

**Figure 1.**
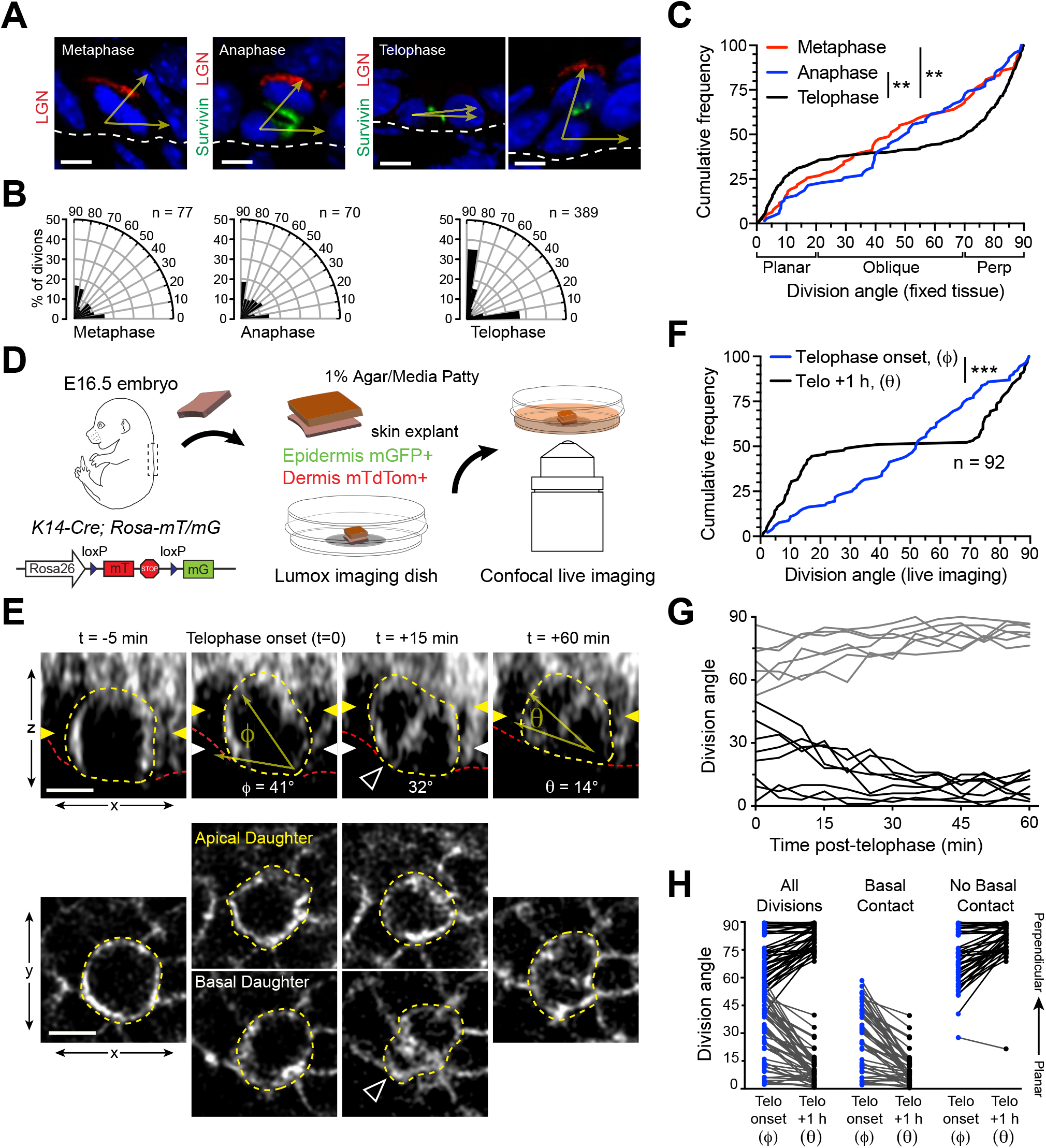
Telophase reorientation corrects oblique anaphase orientations. (**A**) Sagittal sections from E16.5 embryos showing mitotic basal cells at indicated stages. Yellow arrows indicate division axis relative to basement membrane (dashed white line) Apical LGN (red) is generally present in oblique and perpendicular divisions, but absent from planar divisions. Survivin (green) is diffusely distributed between daughter pairs at anaphase, transitioning to stereotypic dual-puncta by telophase. (**B**) Radial histograms of division orientation at metaphase, anaphase, and telophase in E16.5 *wild-type* controls; n indicates number of divisions measured from >20 embryos per mitotic stage. (**C**) Same data as in (**B**), plotted as a cumulative frequency distribution. Note sigmoidal pattern at telophase (black), characteristic of bimodal distribution of division angles. Compare to linear pattern, characteristic of random distributions at metaphase (red) and anaphase (blue). (**D**) Schematic of experimental design for live imaging of embryonic epidermal explants. *K14-Cre; Rosa-mT/mG* is used to label epidermis with membrane (m)-GFP and other tissues (including dermis) with mTdTomato. (**E**) (Top) z-projection stills from a movie of a mitotic cell as it enters anaphase (defined as t=0), through 60 minutes post-anaphase onset, depicting planar telophase correction. Epidermal-dermal boundary shown by red line. Dividing daughter pairs are outlined with yellow dashed lines. Division orientation angles are shown below (ϕ, telophase onset; θ, +1 h). (Bottom), xz en-face views at same time points. Yellow and white arrowheads indicate plane of optical section for apical and basal daughters, respectively. In most cases, planar reorientation is preceded by maintenance of basement membrane contact (open arrowheads), which are most apparent in the en face basal focal plane, where they appear as small membrane circles. (**F**) Cumulative frequency distribution of division angles from live imaging experiments of E16.5 embryos at telophase onset (blue; ϕ) and one hour later (black; θ). n indicates number of divisions from 4 embryos across 4 independent sessions. (**G**) Traces of division orientation at five minute intervals for 15 representative cells from telophase onset to +1 h. Most telophase reorientation generally occurs within ~30 minutes after anaphase onset. (**H**) Data from (**F**) depicting division orientations at telophase and 1 h later. Connecting lines demonstrate that ~60% oblique anaphase divisions reorient to planar (black lines) while the remaining ~40% correct to perpendicular (grey lines). The direction of correction correlates with the maintenance of basal contact following membrane ingression. Scale bars, 5 μm (a), 10 μm (e). ** *P* < 0.01, *** *P* < 0.001, by Kolmogorov-Smirnov test.

Because metaphase and anaphase events are rare in fixed sections, we performed *ex vivo* live imaging of embryonic epidermal explants (Cetera, Leybova, Joyce, & Devenport, 2018), in order to examine the dynamics of spindle orientation at late stages of mitosis (Fig. 1D). We utilized embryos expressing both *ROSA-mT/mG* and *K14-Cre*, where ubiquitous membrane-TdTomato (mT) is recombined to membrane-GFP (mG) specifically in the epidermis. This membrane marker enabled us to visualize the initiation of cleavage furrow ingression, and determine the axis of division at telophase onset (t=0; ϕ=division angle; Fig. 1E).

Importantly, we observed many basal cells with either planar or perpendicular orientations at telophase onset that remained fixed for the duration of the imaging period (Supplementary Fig. 1A,B; Supplementary Videos 1,2). In addition, as suggested by our analyses of fixed tissue, basal progenitors also frequently initiated telophase at oblique angles (Fig. 1E,F). However, these oblique divisions invariably corrected to planar or perpendicular within an hour (Fig. 1E-G; Supplementary Fig. 1C,D; Supplementary Videos 3,4). This reorientation, hereafter referred to as telophase correction, generally occurred within the first 30 minutes after telophase onset. (Fig. 1G). Since little or no reorientation occurred after 1h, we assigned t=+60min as the imaging endpoint (θ=division angle).

While initial orientations of >60° typically corrected to perpendicular, and those <40° to planar, the behavior of intermediate orientations (40-60°) was less predictable (Fig. 1H). However, we noted that apical daughters undergoing planar telophase correction frequently displayed a unique, balloon-shaped morphology and appeared to maintain contact with the basement membrane (open arrowheads in Fig. 1E). Remarkably, maintenance of basal contact predicted planar orientation, while loss of contact predicted the opposite (Fig. 1H). These data suggest that transient oblique metaphase-anaphase orientations are corrected in a manner dependent on whether they retain contact with the basement membrane following cleavage furrow ingression.

### LGN mediates perpendicular telophase correction

Previous studies have shown that LGN (Pins in Drosophila)—along with its binding partners Insc (Inscuteable), NuMA (Mud), and Gαi—play key roles in OCD (Bowman, Neumuller, Novatchkova, Du, & Knoblich, 2006; Du & Macara, 2004; Izumi, Ohta, Hisata, Raabe, & Matsuzaki, 2006; Kraut, Chia, Jan, Jan, & Knoblich, 1996; Mora-Bermudez, Matsuzaki, & Huttner, 2014; Schaefer, Shevchenko, Shevchenko, & Knoblich, 2000; Siller et al., 2006; Williams et al., 2014; Zigman et al., 2005). In the conventional view, LGN functions primarily during prometaphase-metaphase by facilitating capture and anchoring of astral microtubules to the cell cortex. In developing stratified epithelia, LGN first localizes to the apical cortex during prophase (Byrd et al., 2016; Lechler & Fuchs, 2005; Williams et al., 2011; Williams et al., 2014). However, our finding that a large proportion of anaphase cells are oriented obliquely suggests that initial perpendicular spindle positioning by LGN may be imprecise, and raises the question of whether LGN may also function during telophase correction.

We first examined LGN function in fixed tissue using *in utero* lentiviral RNAi with a previously validated shRNA (Williams et al., 2011; Williams et al., 2014). As expected, *LGN*^*1617*^ knockdown shifts the distribution of division angles towards a predominantly planar phenotype. Conversely, LGN overexpression increases the frequency of perpendicular divisions (Supplementary Fig. 2A). This demonstrates that altering LGN levels in either direction impacts division orientation, but does not address whether LGN acts during initial spindle positioning, telophase correction, or both.

Next, we performed live imaging of mosaically-transduced embryonic explants (Fig. 2A). In these experiments, expression of an H2B-RFP reporter allowed us to distinguish transduced/knockdown (RFP+) clones from wild-type internal controls (RFP-). Moreover, this approach enabled tracking of nuclei, facilitating identification of anaphase onset. In control explants transduced with a non-targeting *Scramble* shRNA, similar patterns of telophase correction were observed as in wild-type controls (Fig. 2B), demonstrating that lentiviral transduction has no effect on its own.

**Figure 2.**
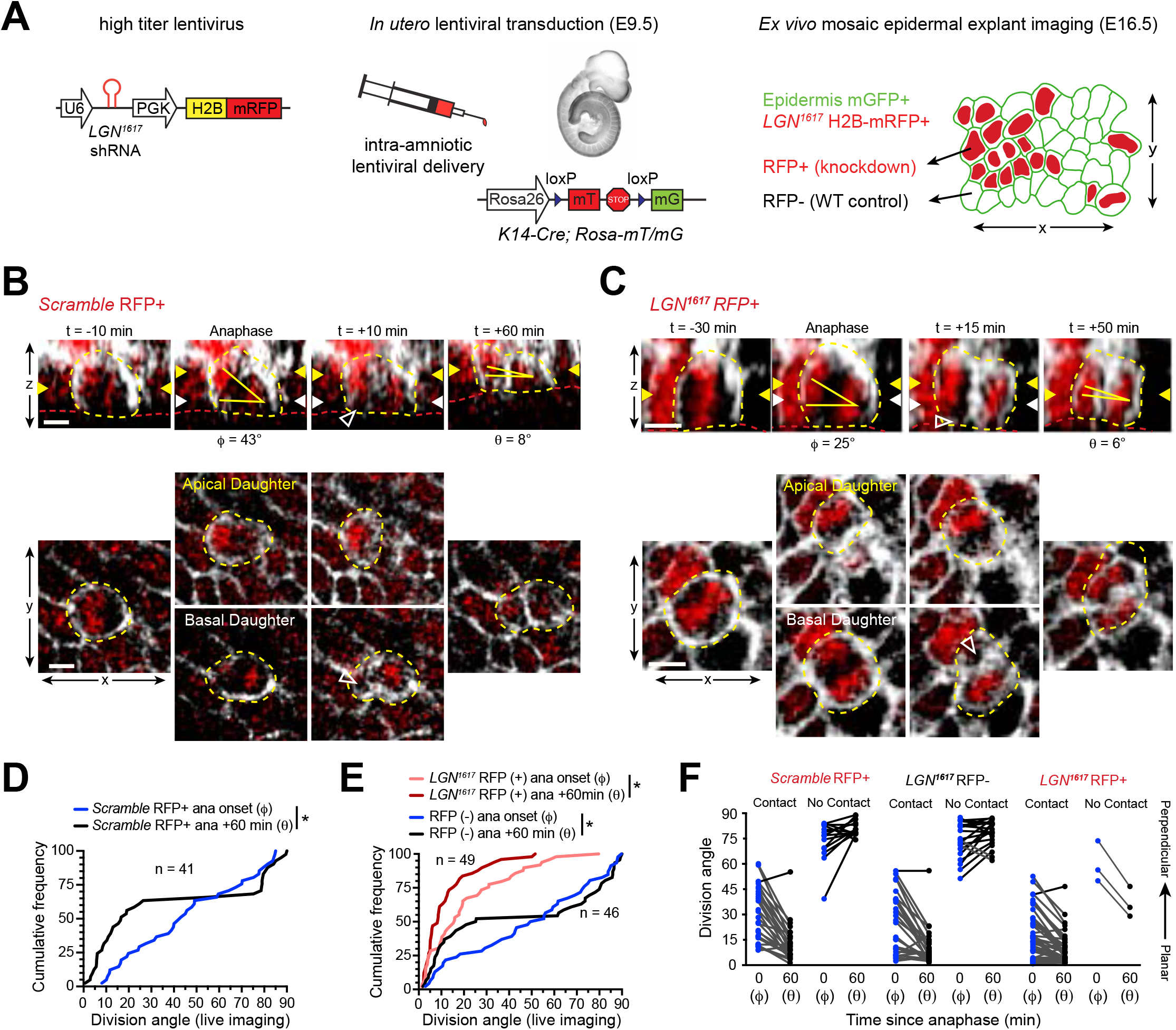
LGN mediates perpendicular but not planar telophase reorientation. (**A**) Schematic of modified experimental protocol of live imaging of epidermal explants (see Fig. 1D) incorporating lentiviral shRNA transduction to generate mosaic knockdown tissue. Transduced/knockdown regions are marked with histone H2B-mRFP1 (H2B-RFP). (**B-C**) Stills from live imaging of (**B**) *Scramble* or (**C**) *LGN*^*1617*^ H2B-RFP+ celsl undergoing planar correction, annotated as in Fig. 1E. Basal contact of obliquely dividing daughter cell is visible in the basal focal plane (open arrowhead). (**D-E**) Cumulative frequency distributions of division orientation from (**D**) *Scramble* or (**E**) *LGN*^*1617*^ H2B-RFP (+/−) live imaging experiments at anaphase onset (ϕ) and 1 h later (θ). *Scramble* RFP-positive and *LGN*^*1617*^ RFP-negative cells display similar patterns of telophase correction as observed in wild-type explants (Fig. 1F). While *LGN*^*1617*^ RFP+ cells are more biased toward planar/oblique at anaphase onset, significant planar correction still occurs; *n* indicates observed divisions from 5 embryos imaged in 4 technical replicates. (**F**) Data from (**D,E**) depicting orientation at anaphase onset (ϕ) and 1 h later (θ) for *Scramble* RFP+ and *LGN*^*1617*^ RFP-negative and RFP+ cells, grouped by presence/absence of basal contact. ~95% of *LGN* knockdown cells correct to planar (θ<30°) 1 h later, though nearly all mitoses maintained basal contact for both daughter pairs. Scale bars, 10μm. * *P* < 0.05 by Kolmogorov-Smirnov test.

Unexpectedly, many *LGN*^*1617*^ RFP+ cells entered anaphase at oblique angles (Fig. 2C). However, while *Scramble* and *LGN*^*1617*^ RFP-negative cells corrected to either planar or perpendicular within 1h of anaphase onset, *LGN*^*1617*^ RFP+ cells invariably corrected towards planar (Fig. 2D,E), suggesting that planar-directed telophase correction is LGN-independent. While nearly all LGN knockdown cells that enter anaphase at oblique angles maintain basal contact, a select few lose contact but still demonstrate planar telophase correction (Fig. 2F; Supplementary Fig. 2B,C). This suggests that, in addition to initial spindle positioning during prometaphase, LGN is required for perpendicular correction during telophase.

### The adhesion protein, vinculin, is essential for telophase correction

Our data suggest that known intrinsic spindle orientation regulators play no role in planar telophase correction. Previous studies have shown that perturbations in epidermal cell-cell adhesions and cell-matrix adhesions can lead to defects in OCD (Dor-On et al., 2017; Lechler & Fuchs, 2005). Coupled with our observation of basal contacts in oblique anaphase divisions, we next decided to probe whether cell-cell or cell-matrix adhesions were required for telophase correction by knockdown of the adherens-junction (AJ) and focal-adhesion actin-scaffold, vinculin (Burridge & Feramisco, 1982; Geiger, 1979). In support of this hypothesis, mosaic knockdown with two distinct shRNAs targeting vinculin (*Vcl*) resulted in randomized telophase division orientation (Fig. 3A; Supplementary Fig. 3A-C). Live imaging of *Vcl*^*3466*^ H2B-RFP explants revealed that vinculin loss had no effect on initial anaphase orientation, but specifically disrupted telophase correction (Fig. 3B,C). While apical daughters were able to retain basal contact upon vinculin loss, *Vcl*^*3466*^ cells no longer corrected in a contact-dependent manner, instead displaying minimal or misdirected radial change (Fig. 3D,E; Supplementary Fig. 3D). These data suggest that proper telophase correction is dependent on cell-cell and/or cell-matrix adhesion.

**Figure 3.**
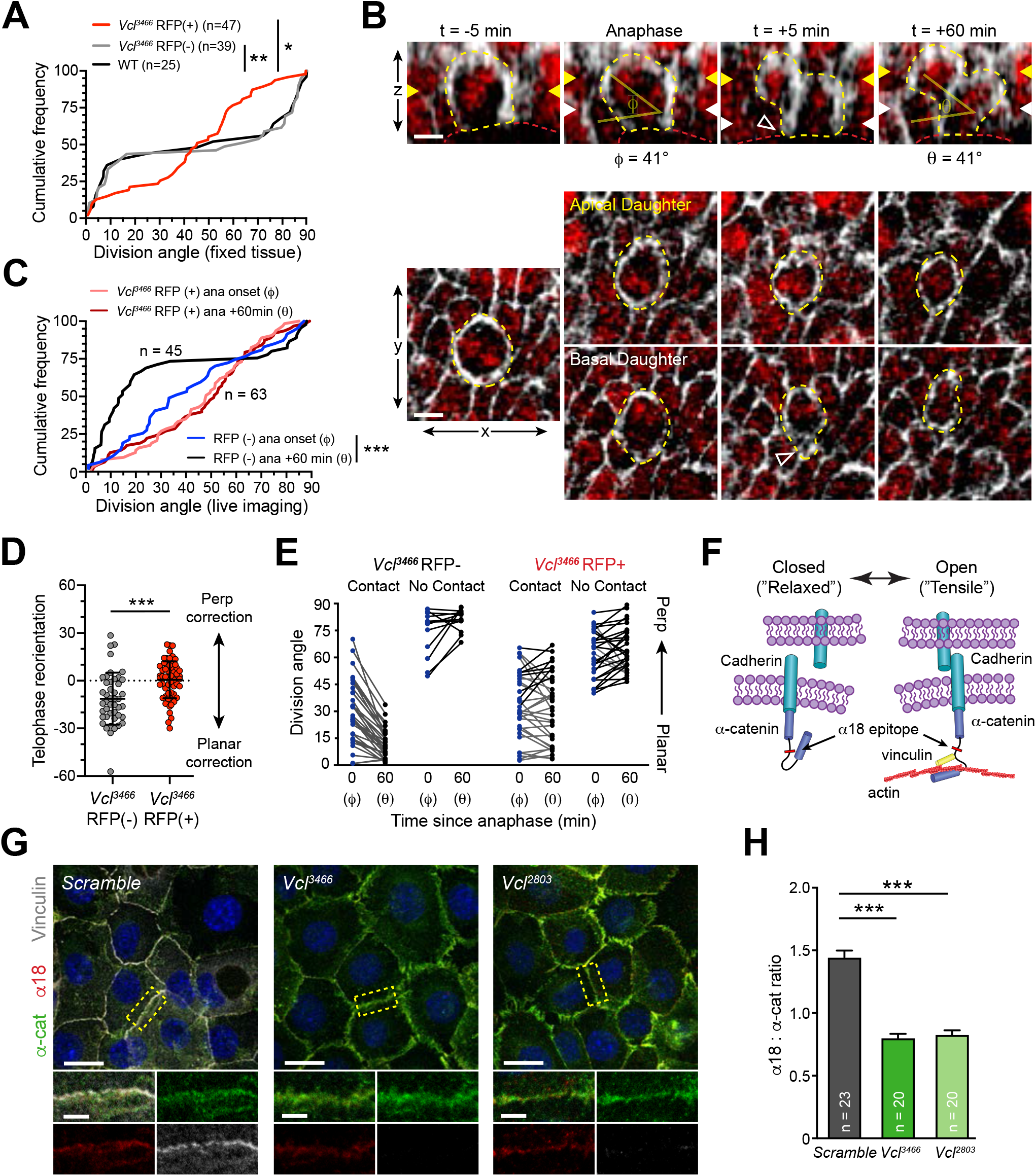
The cell-adhesion/cytoskeletal scaffolding protein vinculin is required for telophase reorientation. (**A**) Cumulative frequency distribution of telophase division angles in *Vcl*^*3466*^ H2B-RFP+ (red), RFP-negative (grey), and littermate controls (WT; black); n indicates number of divisions observed from 3-4 independent embryos. Note bimodal distribution observed in wild-type littermates and RFP-negative cells, in contrast to random distribution of *Vcl*^*2803*^ RFP+ cells. *Vcl*^*2803*^ RFP+ cells (Supplementary Fig. 3B) show a similar phenotype. (**B**) Movie stills of *Vcl*^*3466*^ RFP+ mitotic cell, annotated as in Fig. 1e. While the presence of basal contact (open arrowhead) would predict planar correction, this division remains oblique when reevaluated 1 h later. (**C**) Cumulative frequency distribution of division orientation at anaphase onset (ϕ) and 1 h post-anaphase (θ) for RFP+ and RFP-negative populations, from movies of *Vcl*^*3466*^ mosaic tissue; n indicates divisions from 4 embryos imaged in 3 separate sessions. (**D**) Quantification of telophase reorientation (θ - ϕ) for *Vcl*^*2803*^ RFP+ cells and RFP-controls. (**E**) Data from (**C**) depicting orientation at anaphase onset (ϕ) and 1 h later (θ) for RFP-negative and RFP+ cells. RFP-negative controls sort anaphase orientation (ϕ) into bimodal distribution within 1 h (θ) in a basal-contact dependent manner; *Vcl*^*3466*^ RFP+ cells display minimal change, or correct irrespective of basal contact. (**F**) Cartoon representation of tension-sensitivity model of AJ assembly. In the absence of tension, α-catenin exists in an autoinhibited closed conformation, masking the α18 epitope. In the presence of actin-mediated tension, α-catenin opens, exposing the α18 epitope and vinculin binding domain. (**G**) Stable primary murine keratinocytes cell lines grown in the presence of high (1.5 mM) Ca^2+^ for 8 h form nascent cell-cell adhesions, stained for total α-catenin (green); open, “tensile” α-catenin (α18, red); and vinculin (grey). Single junction magnifications (yellow dashed region) shown below, suggest that vinculin knockdown results in a reduced α18:α-catenin ratio, quantified in (**H**). High magnification single channel images of boxed region shown below; n indicates number of junctions evaluated. Scale bars, 10 μm (**B**), 20 μm (**G**). * P < 0.05, ** P < 0.01, *** P < 0.001, determined by Kolmogorov-Smirnov test (**C**), or unpaired two-tailed student’s t-test (**H**).

In order to differentiate between vinculin’s role in cell-cell and cell-matrix adhesion, we decided to interrogate the function of cell-cell adhesion, specifically. The AJ is canonically composed of transmembrane cadherins, which couple neighboring cells through trans-dimerization in the extracellular space and link to the underlying actin-cytoskeleton via α/β-catenin (Ratheesh & Yap, 2012). In the presence of actin-dependent tension, α-catenin undergoes a conformational change, exposing the α18 epitope (Buckley et al., 2014; Hansen et al., 2013; Rubsam et al., 2017; Yonemura, Wada, Watanabe, Nagafuchi, & Shibata, 2010) (Fig. 3F). Vinculin, itself a tension-sensitive molecule, binds to α-catenin via the internal M-domain and is believed to strengthen the actin binding potential of the AJ (H. J. Choi et al., 2012; Huang, Bax, Buckley, Weis, & Dunn, 2017; Weiss, Kroemker, Rudiger, Jockusch, & Rudiger, 1998).

To probe the functionality of this model in our system, we utilized a calcium-shift adhesion assay in primary cultured keratinocytes (Vasioukhin, Bauer, Yin, & Fuchs, 2000). Following 8-hours of exposure to 1.5mM Ca^2+^, *Scramble* control keratinocytes form linear AJs containing both vinculin and α-catenin (Fig. 3G). While α-catenin is still recruited to AJs in *Vcl* knockdown keratinocytes, in agreement with a recent report (Rubsam et al., 2017), their junctions appeared wider than controls. Loss of vinculin also led to a reduced fluorescence intensity ratio of α18 (‘tensile’) to total α-catenin (Fig. 3H), confirming that the tension sensitivity of α-catenin is vinculin-dependent in keratinocytes.

### Perturbation of AJ components prevents telophase reoriention without affecting LGN

The reduction in tensile α-catenin at AJs observed in *Vcl* mutants—together with the finding that epidermal α-catenin (*Ctnna1*) knockouts display randomized division orientation in early mitosis (Lechler & Fuchs, 2005)—prompted us to ask whether α-catenin might also play a role in telophase correction. Using a previously validated shRNA (Beronja, Livshits, Williams, & Fuchs, 2010), we observed randomized telophase orientation in E16.5 fixed samples upon *Ctnna1* knockdown (Supplementary Fig. 4A,B). Live imaging of mosaic knockdown explants confirmed that α-catenin loss disrupts telophase correction, with no effect on initial anaphase orientation (ϕ; Fig. 4A). Similar to *Vcl* knockdown, *Ctnna1* knockdown disrupted telophase correction, and eliminated the predictiveness of basal contacts for correction directionality (Fig. 4B; Supplementary Fig. 4C). While previous studies have shown that E-cadherin is capable of regulating division orientation through a direct interaction with LGN (Gloerich, Bianchini, Siemers, Cohen, & Nelson, 2017; Hart et al., 2017), knockdown of α-catenin or vinculin produced no measurable defects in LGN recruitment or localization (Fig. 4C-E). These data suggest that proper coordination of cell-cell adhesion and connectivity to the underlying actin cytoskeleton are required for telophase correction.

**Figure 4.**
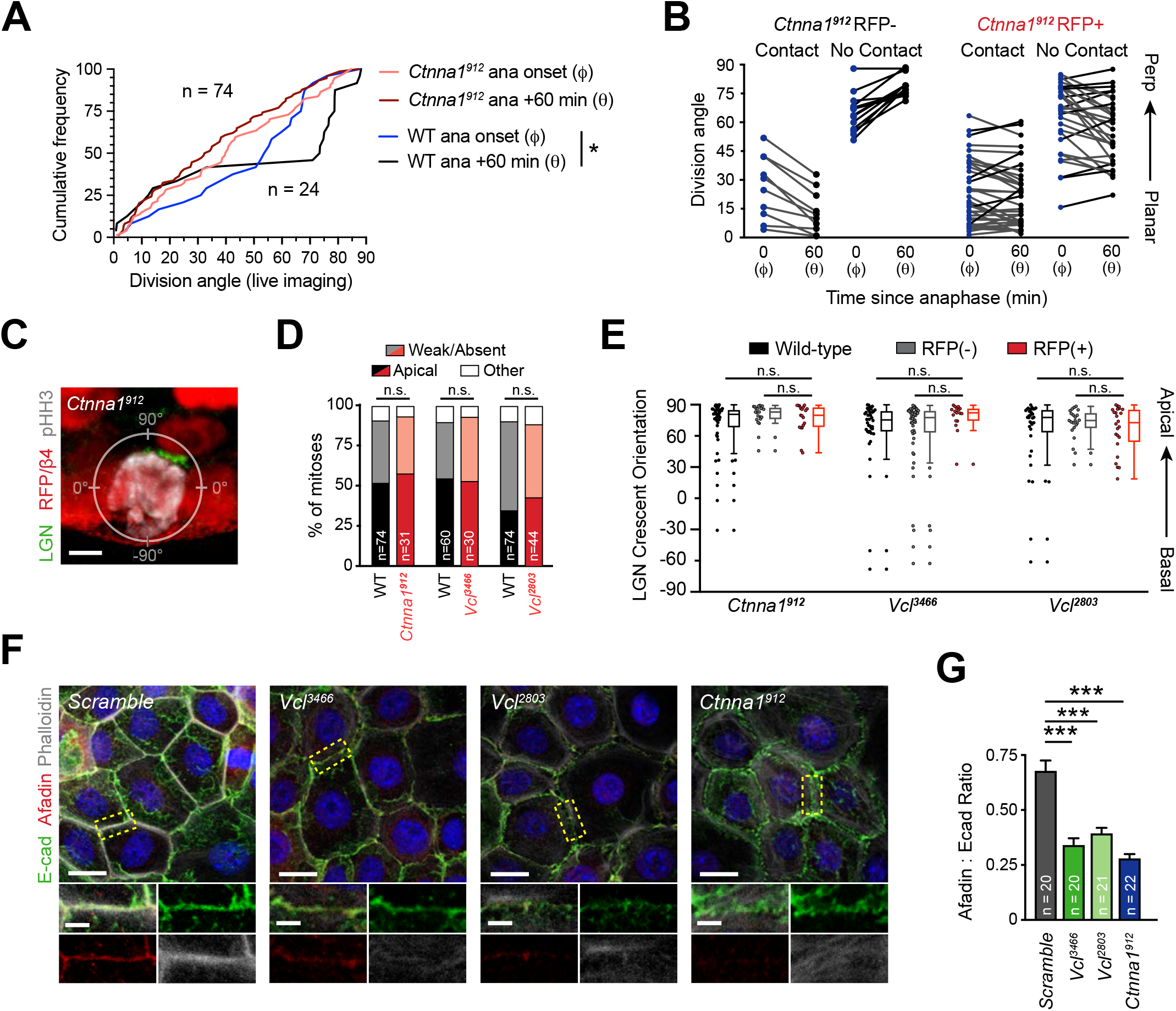
The vinculin/α-catenin cytoskeletal scaffold is essential for telophase reorientation and afadin accumulation in AJs. (**A**) Cumulative frequency distribution of division angles from E16.5 live imaging experiments of *Ctnna1*^*912*^ RFP+ and wild-type littermates; *n* values indicate cells imaged for each group. (**B**) Division orientation at anaphase onset (ϕ) and one hour later (θ) for *Ctnna1*^*912*^ knockdown and wild-type littermates, plotted from data in (**A**). *Ctnna1*^*912*^ RFP+ cells show no obvious correction pattern. (**C**) E16.5 *Ctnna1*^*912*^ RFP+ mitotic cell (pHH3+, white), showing normal apical localization of LGN (green). (**D**) Quantification of LGN recruitment patterns in pHH3+ cells from E16.5 *Ctnna1*^*912*^, *Vcl^3466^*, and *Vcl*^*2803*^ embryos (RFP+ cells), with age-matched littermate controls for each cohort. (**E**) Orientation of LGN crescents from indicated groups. Knockdown of Vcl or Ctnna1 does not significantly alter the tendency of LGN to localize apically. (**F**) Primary mouse keratinocytes after 8 h Ca^2+^ shift—labeled with afadin (red), E-cadherin (green) and actin (phalloidin, grey)—which accumulate in linear bands at cell-cell junctions in Scramble control cells. Yellow boxed region shown at high magnification below; n indicates junctions evaluated. Vcl and Ctnna1 knockdown cells show defects in linear actin accumulation as well as afadin recruitment to AJs, quantified in (**G**). Scale bars, 5μm (**C**), 20μm (**F**). P values determined by Kolmogorov-Smirnov test (**A**), χ^2^ (**D**), or student’s unpaired t-test (**E,G**). * P < 0.05, *** P < 0.001. n.s. not significant (P > 0.05).

### Vinculin and α-catenin are required for afadin recruitment to nascent AJs

Previous studies have demonstrated that the actin scaffold afadin is capable of binding directly to α-catenin via an internal domain proximal to the vinculin binding domain (Mandai et al., 1997; Pokutta, Drees, Takai, Nelson, & Weis, 2002). This prompted us to address whether afadin junctional recruitment is regulated by the α-catenin/vinculin complex. In calcium-shift assays, afadin demonstrated robust coaccumulation with E-cadherin after 8 hours in *Scramble* controls (Fig. 4F). However, junctional afadin recruitment was significantly reduced by knockdown of vinculin or α-catenin (Fig. 4F,G). This coincided with reduced actin accumulation and altered E-cadherin continuity (Supplementary Fig. 4D), such that AJs displayed a persistent punctate or ‘zipper’ morphology upon vinculin or α-catenin loss (Vasioukhin et al., 2000). These data reinforce the role of these proteins in adhesion formation, likely related to their actin-binding capabilities.

### Afadin regulates telophase correction and basal contact stability

In Drosophila, the afadin homologue Canoe (Cno) is essential for asymmetric cell division of embryonic neuroblasts (Speicher, Fischer, Knoblich, & Carmena, 2008). Recent studies in mammals have similarly described a role for afadin in regulating division orientation in the embryonic kidney and cerebral cortex (Gao et al., 2017; Rakotomamonjy et al., 2017). To interrogate the effects of afadin (*Afdn*) loss in the epidermis we utilized three different models of afadin knockdown (*Afdn*^*2711*^) or conditional knockout (*Afdn*^*fl*/*fl*^, crossed to *K14-Cre* or injected with lentiviral Cre-RFP) (Supplementary Fig. 5A,B). Each model displayed randomized telophase orientation in E16.5 fixed sections and whole-mounts (Fig. 5A; Supplementary Fig. 5C,D). Results from live imaging experiments showed afadin loss has no effect on anaphase orientation, while oblique divisions fail to undergo directed telophase correction (Fig. 5B,C; Supplementary Fig 5E; Supplementary Video 5). While *Scramble* controls and LGN mutants demonstrated predominantly planar telophase correction, *Afdn* knockdown phenocopied loss of vinculin and α-catenin, with minimal or randomized reorientation of oblique divisions, particularly when basal contact is not maintained (Fig. 5D; Supplementary Fig. 5F).

**Figure 5.**
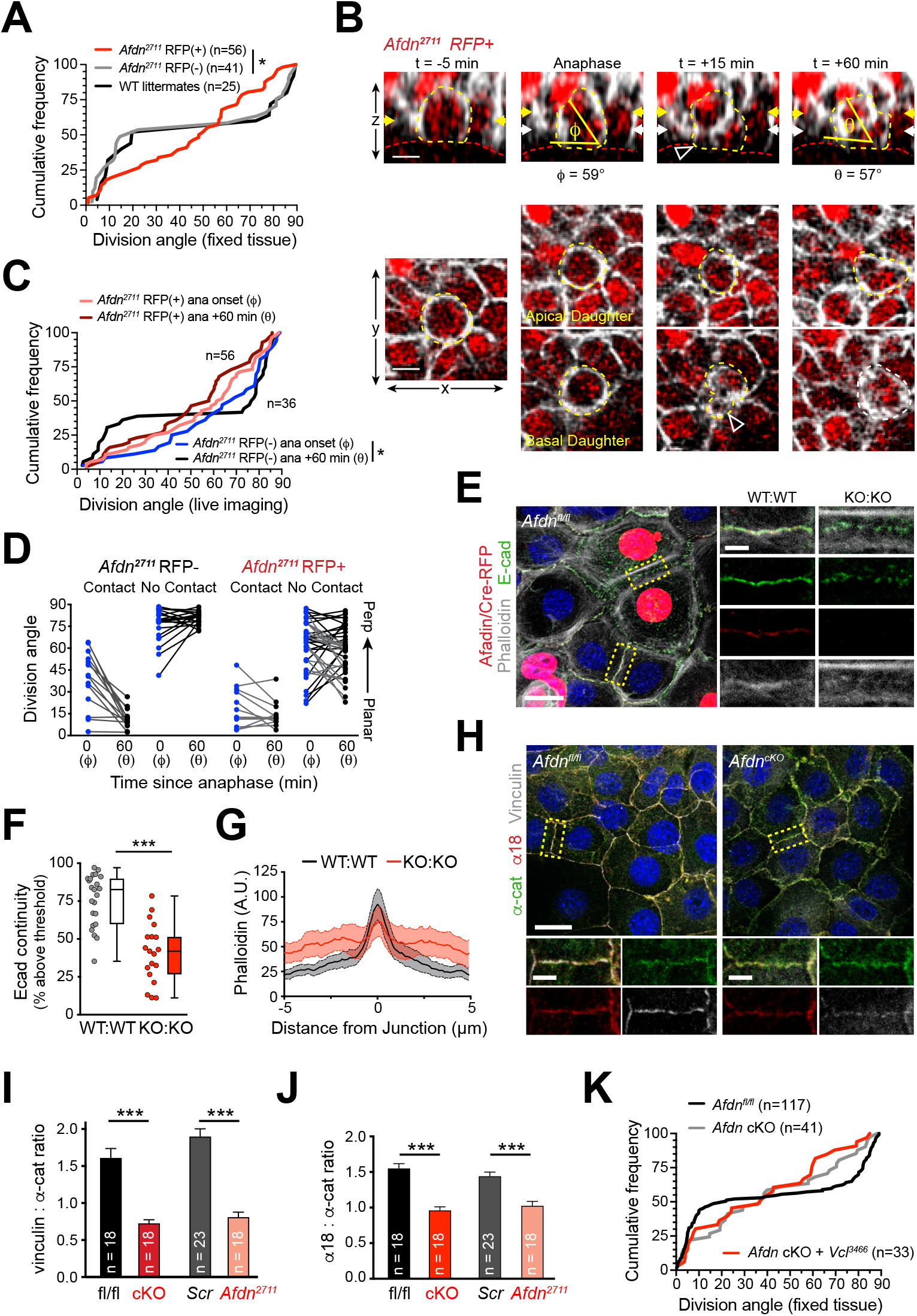
Afadin is essential for telophase correction and basal contact maintenance. **(A)** Cumulative frequency distribution of telophase division angles from E16.5 epidermis. Afdn knockdown (red) results in random telophase orientation, while uninfected RFP-negative (grey) and littermate (black) controls display a bimodal distribution; n indicates cells analyzed from 3-6 embryos. **(B-E)** Live imaging of E16.5 *Afdn*^*2711*^ explants. **(B)** An obliquely-oriented *Afdn*^*2711*^ RFP+ cell fails to reorient, while losing basal contact (open arrowhead). **(C)** Cumulative frequency distributions of division orientation from E16.5 live imaging of *Afdn*^*2711*^ RFP+ and wild-type littermates; n indicates observed divisions from 3 embryos imaged in 2 separate sessions. **(D)** Division orientation at anaphase onset (ϕ) and one hour later (θ) for *Afdn*^*2711*^ RFP+ and RFP-negative cells, plotted from data in **(C)**. RFP-negative controls correct into a bimodal distribution, while RFP+ cells reorient randomly. **(E)** *Afdn*^*fl*/*fl*^ primary keratinocytes mosaically infected with Cre-RFP (red) after 8 h 1.5 mM Ca^2+^ shift, stained for E-cadherin (green), afadin (red), and phalloidin (grey). Junctions between two uninfected cells (WT:WT) show linear morphology with consistent E-cadherin (green), afadin (red) and phalloidin (grey) labeling. In contrast, junctions between two infected cells are punctate, with less junction-associated phalloidin. **(F)** Quantification of E-cad continuity along junction length, as in Fig. 4J. **(G)** Quantification of fluorescence intensity of actin (phalloidin) measured by orthogonal linescans. Phalloidin is decentralized in KO:KO junctions (red) compared to WT:WT (black;. n indicates junctions evaluated. **(H)** Primary keratinocytes derived from *Afdn*^*fl*/*fl*^ K14-Cre+ embryos (*Afdn*^*cKO*^) and Cre-negative littermates after 8 h Ca^2+^ shift and stained for α18 (red), α-E-catenin (green), and vinculin (grey). **(I)** Quantification of vinculin:total α-catenin and **(J)** α18:total α-catenin fluorescent ratios; n indicates junctions analyzed. **(K)** Cumulative frequency distribution of E16.5 telophase division angles in *Afdn*^*fl*/*fl*^, *Afdn*^*cKO*^, and *Afdn*^*cKO*^ + *Vcl*^*3466*^ H2B-RFP epidermis. *Vinculin* knockdown does not exacerbate Afdn knockout phenotype. Scale bars, 10μm **(B)**, 20μm **(E,H)**. P values determined by Kolmogorov-Smirnov test **(A,C,K)**, student’s unpaired t-test **(F,I,J)**. * P < 0.05, *** P < 0.001.

### Afadin is required for normal AJ morphology and is a novel regulator of α-catenin conformation

Afadin and Cno are required to stabilize actin-AJ associations during moments of high actomyosin contractility, suggesting a role in establishing/maintaining tensile loads (W. Choi et al., 2016; Sawyer et al., 2011). To examine whether afadin loss influences AJ-associated actin in keratinocytes, we generated mosaic cultures of wild-type and *Afdn*^*cKO*^ cells by transducing *Afdn*^*fl*/*fl*^ keratinocytes with lentiviral Cre-RFP (Fig. 5E). E-cadherin^+^ AJs between wild-type uninfected cells (WT:WT) showed normal accumulation of afadin, while junctions between RFP+ cells (KO:KO) lacked afadin (Fig. 5E, red). *Afdn*^*cKO*^ cells also demonstrated increased levels of cytoplasmic E-cadherin, and KO:KO junctions displayed punctate, rather than linear, E-cadherin (Fig. 5F), reminiscent of immature “spot” junctions seen after 30 minutes of Ca^2+^ exposure (Supplementary Fig. 5G-I). In addition, while WT:WT junctions showed tight association of actin with E-cad, KO:KO junctions showed reduced junctional actin, with actin bundles frequently displaced ~1 μm from the junction (Fig. 5E,G). Similar results were observed when comparing homogenous cultures of *Scramble* control and *Afdn*^*2711*^ knockdown keratinocytes (Supplementary Fig. 5J-L). These data suggest that afadin plays an essential role in linking cortical actin to the AJ complex, with potential consequences on E-cadherin clustering.

Since it has been shown that AJ components such as E-cadherin regulate junctional recruitment of vinculin from focal adhesions in a tension dependent manner (Noethel et al., 2018; Rubsam et al., 2017), and we noted that α-catenin and vinculin are required for afadin accumulation in the AJ, we wondered whether afadin reciprocally regulates α-catenin or vinculin. Knockout or knockdown of *Afdn* resulted in decreased junctional accumulation of both vinculin and the α18 epitope, reducing both vinculin:α-catenin and α18:α-catenin fluorescence intensity ratios (Fig. 5I,J). This suggests that afadin is a novel regulator of AJ tension sensitivity by affecting α-catenin conformation and vinculin recruitment.

Finally, we sought to test genetically whether afadin and vinculin operate in the same molecular pathway in the context of telophase correction. To do so, we performed embryonic lentiviral injection of vinculin shRNA on an *Afdn*^*cKO*^ or *Afdn*^*fl*/*fl*^ background. Examination of division orientation in single and double mutants revealed that additional loss of vinculin did not exacerbate the *Afdn*^*cKO*^ phenotype, suggesting these proteins operate in a linear pathway in the context of division orientation (Fig. 5K).

### Telophase correction occurs independently of canonical polarity and spindle-orienting cues

In *Drosophila*, Cno is essential for early establishment of apical-basal polarity during cellularization (Bonello, Perez-Vale, Sumigray, & Peifer, 2018; W. Choi, Harris, Sumigray, & Peifer, 2013). A similar role has been described for afadin in mammalian development (Komura et al., 2008; Rakotomamonjy et al., 2017; Yang et al., 2013). Furthermore, both Par3 and its Drosophila ortholog Bazooka are required for OCD via regulation of LGN localization (Schober, Schaefer, & Knoblich, 1999; Williams et al., 2014; Wodarz, Ramrath, Kuchinke, & Knust, 1999). Thus, we asked how afadin loss impacts expression of the canonical apical polarity cue Par3. In controls, Par3 accumulates at the apical cortex throughout the cell cycle (Supplementary Fig. 6A). We measured Par3 radial fluorescence intensity at interphase and determined that knockout of Afdn reduced apical accumulation 15-30% (Supplementary Fig. 6B). However, this had no effect on the apical positioning of centrosomes, suggesting that apical-basal polarity remains largely intact in *Afdn* mutants (Supplementary Fig. 6C,D).

Previous studies in *Drosophila* have shown that Cno interacts directly with Pins and regulates its cortical recruitment (Speicher et al., 2008; Wee, Johnston, Prehoda, & Doe, 2011), while the mammalian orthologs can also directly interact in HeLa cells (Carminati et al., 2016). Therefore, we sought to determine whether a direct interaction between afadin and LGN could underlie the spindle orientation defect observed in *Afdn* mutants. As with *Vcl* and *Ctnna1* mutants (Fig. 4C-E)—and similar to what has been observed in *Drosophila* neuroblasts upon *Cno* loss (Speicher et al., 2008)—*Afdn* loss did not impact the apical recruitment of LGN during early stages of mitosis (Fig. 6A,B; Supplementary Fig. 6E-G). In *Drosophila* neuroblasts, genetic epistasis and protein localization studies support the view that Cno/afadin acts downstream of Pins/LGN and upstream of Mud/NuMA (Speicher et al., 2008). However, we find that neither NuMA, nor its downstream binding partner dynactin, appears to be mislocalized in *Afdn* mutants (Fig. 6C,D; Supplementary Fig. 6H). In addition, NuMA staining overlapped with LGN in early mitotic cells, regardless of afadin presence/absence (91% in Afdnfl/fl, n = 22; 93% in AfdncKO, n = 14).

**Figure 6.**
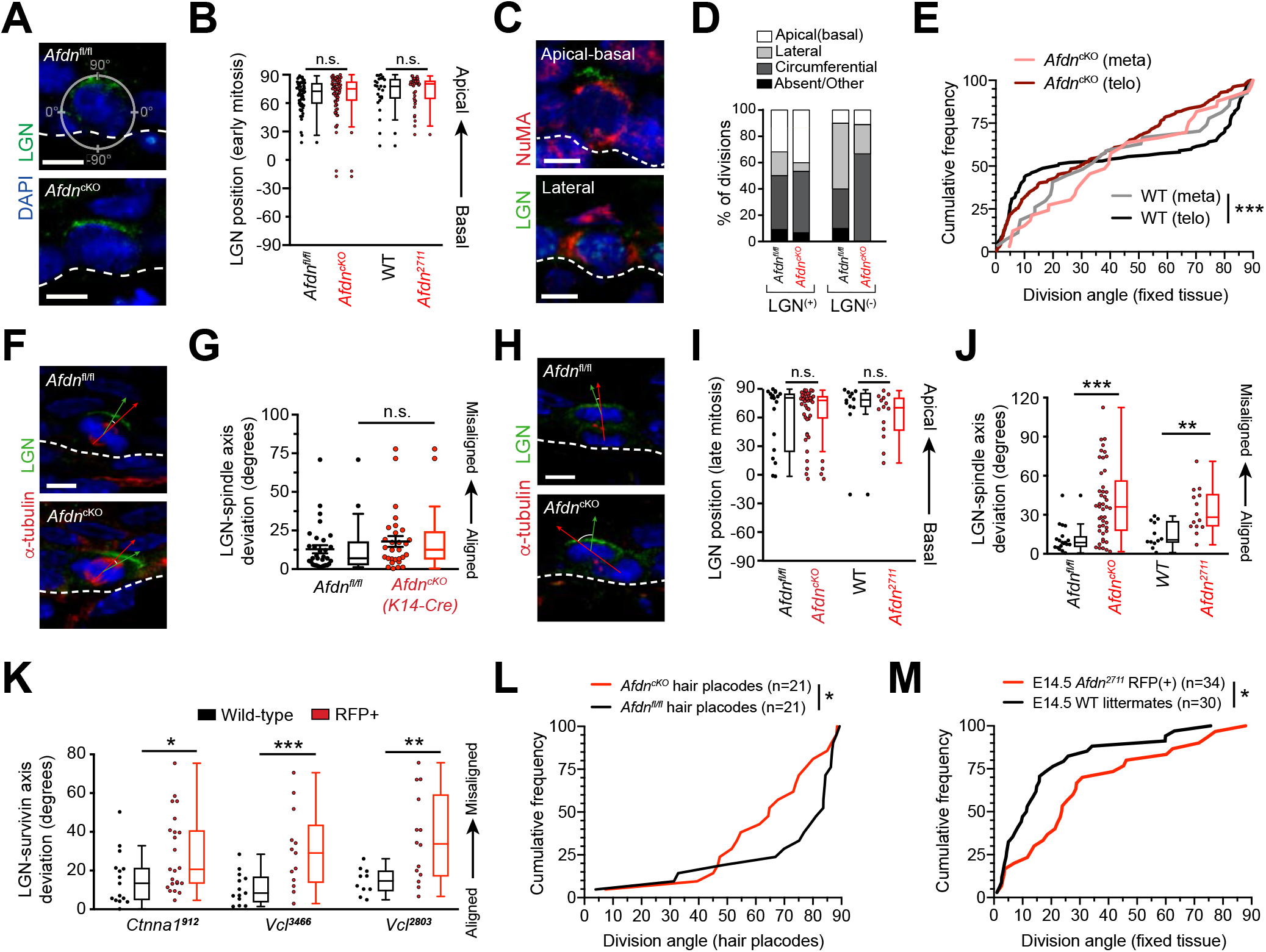
AJ mutants orient their mitotic spindle appropriately but fail to align division plane to LGN in telophase. (**A**) Immunostaining for LGN (green) in E16.5 *Afdn*^*fl*/*fl*^ and *Afdn*^*cKO*^ epidermis. LGN localizes at the apical cortex during mitosis regardless of afadin loss. (**B**) Orientation of LGN crescents in E16.5 mitotic cells from indicated groups. Knockdown or knockout of afadin does not significantly alter the tendency of LGN to localize apically. (**C**) NuMA (red) immunostaining in E16.5 mitotic cells. NuMA localizes predominantly in a bipolar manner, but displays unique patterns in the presence/absence of LGN (green). (**D**) Quantification of NuMA localization patterns binned by LGN presence/absence and genotype. Knockout of *Afdn* does not alter NuMA accumulation. (**E**) Cumulative frequency distribution of spindle orientation at metaphase, and division orientation at telophase, in *Afdn*^*cKO*^ and *Afdn*^*fl*/*fl*^ control littermates. WT spindle orientation (grey line) is random at metaphase before becoming bimodal at telophase (black). In contrast, *Afdn*^*cKO*^ orientation remains random at both metaphase (pink) and telophase (red); n indicates number of observed divisions from 2-3 independent embryos. (**F-H**) Costaining of E16.5 metaphase (**F**) and telophase (**H**) divisions with α-tubulin (red) and LGN (green) in *Afdn*^*cKO*^ and *Afdn*^*fl*/*fl*^ control littermates. (**G**) Quantification of the deviation between the metaphase spindle axis (red arrow in **F**) and LGN radial orientation (green arrow in **F**). Afdn knockout does not disrupt spindle-LGN linkage. (**I**) Quantification of LGN localization in telophase (green arrow in **H**). LGN localizes to the apical cortex of the apical daughter in afadin knockdown/knockout. (**J**) Quantification of the alignment between telophase division orientation (red arrow in **H**) with orientation of LGN (green arrow in **H**). *Afdn*^*cKO*^ cells demonstrate oblique telophase orientation despite normal localization of LGN. (**K**) Deviation between telophase division axis (labeled with survivin) and LGN orientation upon afadin, α-catenin, or vinculin knockdown (as measured in **K**). α-catenin and vinculin loss phenocopies *Afdn* knockout/knockdown. (**L**) Cumulative frequency distribution of division orientation in E16.5 hair placodes (determined by P-cad staining). *Afdn* knockout (red) results in an increase in oblique orientations when compared to predominantly perpendicular orientation observed in *Afdn*^*fl*/*fl*^ control littermates; *n* indicates observed divisions from 3-4 independent embryos. (**M**) Cumulative frequency distribution of E14.5 division orientation in *Afdn*^*fl*/*fl*^ controls (black) and *Afdn*^*cKO*^ (red). Afdn knockout increases the frequency of oblique OCDs; *n* indicates observed divisions from 3-4 independent embryos. Scale bars, 5 μm. P values determined by student’s t-test or Mann-Whitney test depending on tests of normality (**B,G,I-K**), Kolmogorov-Smirnov test (**E,L,M**), or χ^2^ (**D**). * P < 0.05, ** P < 0.01, *** P < 0.001. n.s. not significant (P > 0.05).** P < 0.01.

We previously demonstrated that the mitotic spindle can become misaligned with cortical LGN during metaphase, e.g. following NuMA knockdown (Williams et al., 2011). Thus, we sought to examine whether *Afdn* loss could also lead to uncoupling of the division axis from LGN polarity cues at any point during mitosis, perhaps independently of NuMA. To test this, we co-stained prefixed E16.5 *Afdn*^*fl*/*fl*^ and *Afdn*^*cKO*^ sections with LGN and α-tubulin in order to visualize spindles during metaphase, and cleavage furrow ingression later in telophase. Importantly, while afadin loss altered telophase division orientation, it had no effect during metaphase, where spindles were randomly-oriented (Fig. 6E). Furthermore, the apical LGN crescent aligned with the metaphase spindle axis—regardless of its orientation—in both *Afdn*^*cKO*^ and control embryos (Fig. 6F,G). However, in telophase cells, while LGN remained apically-positioned in both controls and *Afdn* mutants, the orientation of the spindle axis became uncoupled from LGN in *Afdn* mutants (Fig. 6H-J, Supplementary Fig. 6I). Similarly, knockdown of α-catenin or vinculin phenocopied afadin loss, demonstrating that AJ perturbation does not alter LGN localization, but does affect the ability of telophase cells to reorient in response to apical cues (Fig. 6K; Supplementary Fig. 6J).

These findings—together with our observation that afadin and LGN mutants differ in their planar telophase correction phenotypes—suggest that afadin acts independently of LGN in the context of spindle orientation. As further evidence, we find that while LGN strongly colocalizes with known binding partners Gαi3 and Insc, afadin demonstrates minimal colocalization with LGN either pre- or post-chromosome segregation (Supplementary Fig. 6K-M). Finally, there are several contexts during epidermal development where LGN is not required for division orientation. First, although hair placode progenitors undergo perpendicular asymmetric divisions (Ouspenskaia et al., 2016), LGN is weakly expressed in mitotic placode cells, and is not required for proper division orientation (Byrd et al., 2016; Ouspenskaia, Matos, Mertz, Fiore, & Fuchs, 2016). Second, while *LGN* loss reduces perpendicular divisions in the interfollicular epidermis at E16.5, LGN is dispensable at E14.5, when the majority of divisions are planar and LGN is rarely cortical (Williams et al., 2014). In contrast, *Afdn* knockdown increases the frequency of oblique divisions in both contexts, suggesting an LGN-independent function for afadin in both perpendicular and planar divisions (Fig. 6L,M). Together, these data suggest that afadin is a minor or transient LGN-interactor *in vivo* and support a polarity- and LGN-independent role for afadin in telophase correction.

### Telophase correction also occurs during early stratification

The observations that afadin is required for telophase correction at E16.5 (Fig. 5B-D), and that *Afdn* mutants display division orientation defects at early as well as peak stages of stratification (Fig. 5A; Fig. 6M) prompted us to examine whether telophase correction occurs throughout epidermal morphogenesis. To do so, we performed live imaging on wild-type *K14-Cre;mT/mG* epidermal explants E14.5 (Fig. 7A). Even though nearly all divisions at E14.5 are planar (Lechler & Fuchs, 2005), remarkably, at telophase onset, the distribution of observed orientations was randomly-distributed, similar to what was observed at E16.5 (Fig. 7B; compare to Fig. 1F). However, while 47% of cells (n=78) entered telophase oriented obliquely, the vast majority of these maintained apical daughter basal contacts and corrected to planar within 1h of telophase onset (Fig. 7A-C). On the other hand, the few cells (28%) that did not maintain basal contact corrected randomly at E14.5, in contrast to E16.5, when they invariably corrected to perpendicular (compare Fig. 7C to Fig. 1H). Since LGN does not localize cortically or influence division orientation at E14.5 (Williams et al., 2014), this provides additional evidence that LGN is required for perpendicular telophase correction.

**Figure 7.**
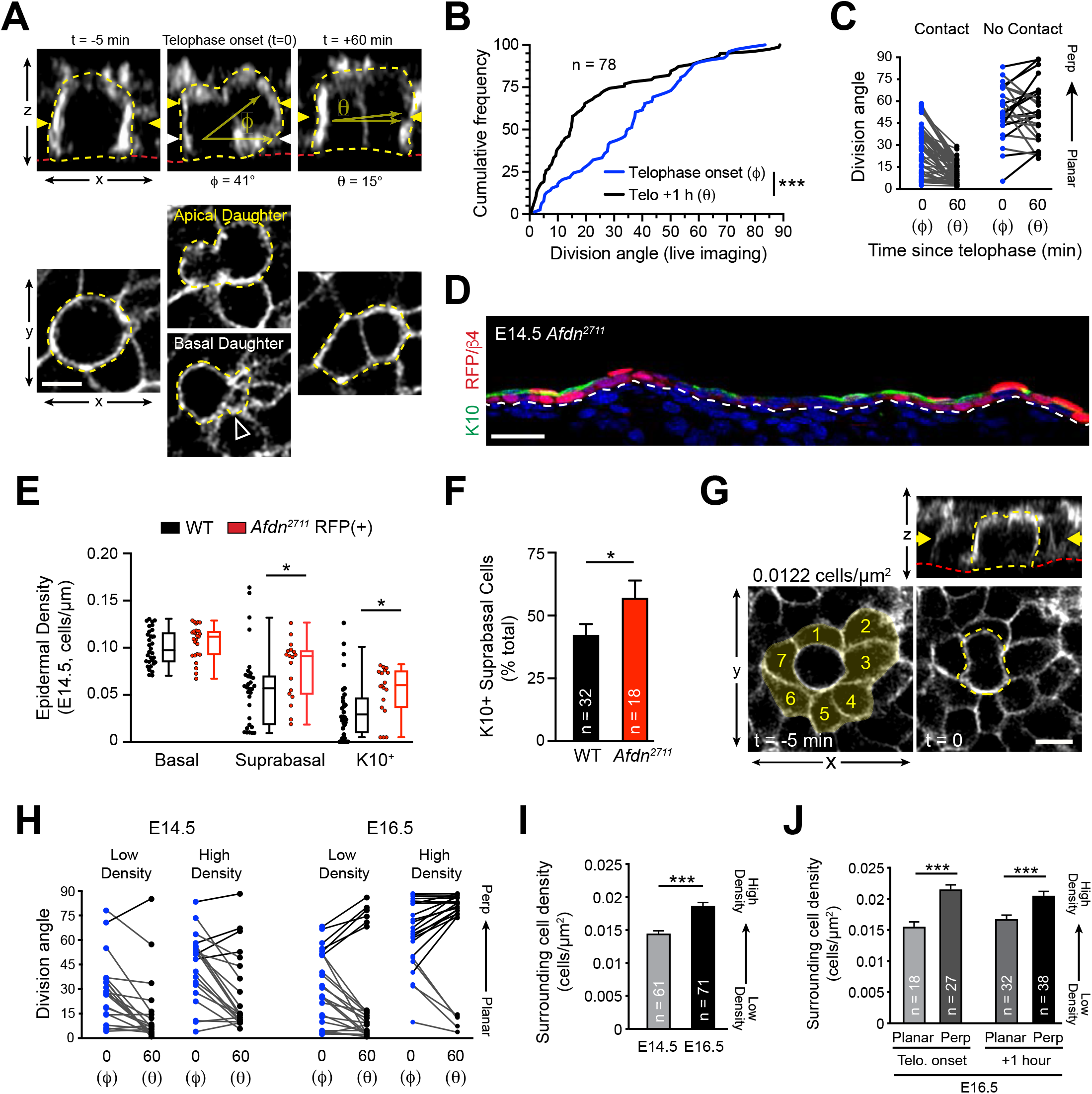
Planar telophase correction limits precocious differentiation in early epidermal development. (**A**) (Top) z-projection stills from a movie of an E14.5 mitotic cell, annotated as in Fig. 1E. (**B**) Cumulative frequency distribution of division angles from live imaging experiments of E14.5 embryos at telophase onset (blue; ϕ) and one hour later (black; θ). n indicates number of divisions from 3 embryos across 2 independent sessions. (**C**) Data from (**B**) depicting division orientations at telophase and 1 h later, sorted based on retention/loss of basal contact throughout cell division. Connecting lines demonstrate that, at E14.5, planar correction occurs in a contact dependent manner, while mitoses that lose contact demonstrate no obvious pattern of correction. (**D**) Sagittal section of E14.5 epidermis with mosaic *Afdn*^*2711*^ H2B RFP transduction. Regions of high infection display increased stratification, as demonstrated by K10 (green) positivity. (**E**) Quantification of epidermal differentiation at E14.5 in WT and *Afdn*^*2711*^ mutants. *Afadin* knockdown increases suprabasal and K10+ cell density, suggesting precocious differentiation. (**F**) Quantification of K10+ suprabasal proportion in E14.5 WT and *Afdn*^*2711*^ mutants. (**G-J**) Relationship between mitotic neighbor cell density and division orientation. n indicates number of mitoses from 3-4 embryos from 2-4 independent imaging sessions. n indicates number of mitoses from 4-6 embryos. (**G**) En-face (bottom) and z-projection (top) stills from movies of WT E14.5 epidermis just before (t = −5 min) and at anaphase onset (t = 0). (**H**) Surrounding cell density (e.g., the area of cells 1-7 in (**G**) at t=-5 min; highlighted in yellow) correlates with distinct division orientation behaviors. Low density mitoses (least dense third of observed divisions) are more likely to display planar anaphase orientation and telophase correction than high density mitoses (most dense third) at both E14.5 and E16.5. (**I**) Comparison of surrounding cell density at E14.5 and E16.5. (**J**) Quantification of surrounding cell density in E16.5 WT epidermis, sorted by telophase onset (ϕ) vs 1-hour later (θ). Planar divisions (0-30°) occur in regions of lower cell density when compared to perpendicular divisions (60-90°). Scale bars, 10μm (**A,G**). P values determined by Kolmogorov-Smirnov test (**B**), student’s unpaired t-test (**E,F,J**), or Mann-Whitney test (**I**). * P < 0.05, *** P < 0.001.

### Afadin loss induces precocious differentiation

At E14.5, *Afdn* mutants displayed an increased proportion of oblique—presumably asymmetric—divisions compared to controls (Fig. 6M), which led us to ask whether afadin loss could promote differentiation. In E14.5 *Afdn*^*2711*^ mosaic epidermis, we noted that Keratin-10 (K10)—a marker of differentiated cells—was enriched in RFP+ mutant regions compared to RFP-negative wild-type regions (Fig. 7D). While progenitor (basal cell) density was similar between *Afdn*^*2711*^ embryos and non-transduced littermates, the density of differentiated cells—whether assessed by their suprabasal position or K10 expression—was significantly higher in *Afdn* mutants (Fig. 7E,F). We conclude that *Afdn* loss induces precocious differentiation, likely through the failure of oblique divisions to correct to planar during telophase.

### Local cell density influences division orientation

Since basal cells become progressively smaller and less elongated between E12.5 and E14.5 (Luxenburg et al., 2015), we wondered whether this this trend continued into later development, and if so, whether increasing cell density could explain the surge in perpendicular divisions that occurs at E16.5. To correlate cell density with division orientation, we analyzed stills from movies of wild-type *K14-Cre; mT/mG* epidermal explants, using the density of mitotic cell neighbors as a measure of local crowding (Fig. 7G). By this metric, mitotic cell neighbor density increased significantly between E14.5 and E16.5 (Fig. 7H). Next, we asked how cell density affected initial telophase division orientation and correction behavior. At E14.5, perpendicular divisions are rare, but most of them occur in high density regions; however, this correlation is much more striking at E16.5 (Fig. 7I). Whether assessing division orientation at either telophase onset or completion, perpendicular orientations consistently occurred in regions of high local density at E16.5 (Fig. 7J). Thus, local cell density not only affects where cells divide (Miroshnikova et al., 2018), but also how they divide, with respect to their division axis.

There is a precedent for AJ components acting as density sensors. For example, epidermal-specific α-catenin knockouts display hyperplasia, likely due to α-catenin-driven negative regulation of Yap1, the transcriptional coactivator of the growth-control Hippo pathway (Schlegelmilch et al., 2011; Vasioukhin, Bauer, Degenstein, Wise, & Fuchs, 2001). Additionally, E-cadherin trans-interactions in MDCK cells are essential to limit both Yap1 and β-catenin transcriptional activation and cell-cycle entry (Benham-Pyle, Pruitt, & Nelson, 2015). Although we did not observe a significant difference in basal cell density between *Afdn* mutants and littermates at E14.5 (Fig. 7E), these studies led us to wonder whether AJ perturbation could affect division orientation by affecting cell crowding and/or density sensing. In agreement with our previous observations in E16.5 wild-type explants, neighbors of control *Scramble* H2B-RFP+ cells with planar anaphase orientations were less crowded when compared to neighbors of cells with perpendicular anaphase orientations (Supplementary Fig. 7A,B). However, while *Afdn* knockdown slightly increased cell density overall, the correlation between local cell density and division orientation persisted (Supplementary Fig. 7C). Thus, while loss of afadin does not appear to perturb density sensing, these data support previous claims that cell crowding can instigate differentiation, with division orientation acting as a potential relief mechanism.

## DISCUSSION

### A two-step mechanism for OCD

These studies shed new light on the mechanisms governing OCDs in the developing epidermis. While previous studies have demonstrated essential roles for canonical spindle orientation genes in division orientation, we now show that initial spindle positioning is only one part of the process. Our data suggests that LGN and associated proteins operate early in mitosis to promote perpendicular divisions, but do so with a high degree of imprecision, resulting in a wide distribution of anaphase division angles between 45-90°. While this function of LGN is required for perpendicular divisions to occur, this fails to explain the bimodal distribution of division angles observed in telophase. In the second phase of our model, telophase cells undergo dynamic reorientation towards a planar or perpendicular orientation, where the direction of correction is dependent on contact with the basement membrane (Fig. 8). This phase relies on the actin-scaffolding α-catenin/vinculin/afadin pathway, highlighting a role for cell adhesion and cytoskeletal dynamics in division orientation. Moreover, our findings now provide an explanation for the randomized division orientation observed in α-catenin mutants more than a decade ago (Lechler & Fuchs, 2005). Lastly, while our data show that LGN is not required for planar-directed telophase reorientation, it suggests that LGN, perhaps in concert with other apical polarity cues, may regulate telophase reorientation in the perpendicular direction, expanding their role in promoting perpendicular divisions into the late stages of mitosis.

**Figure 8.**
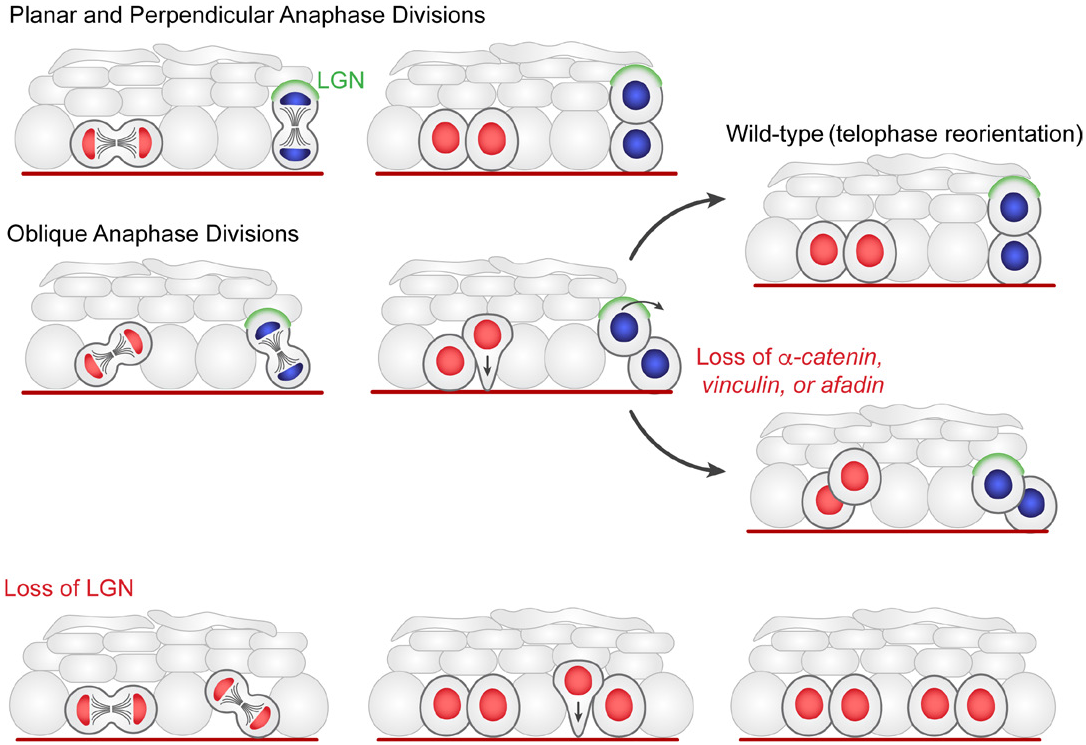
Two-step model of division orientation in the embryonic epidermis. During stratification, LGN (green) is recruited to the apical cortex in ~50% of mitoses, promoting perpendicular divisions. For OCDs with perpendicular and planar anaphase orientations, the division angle is fixed at anaphase onset, exhibiting minimal change in radial orientation during telophase. Importantly, the activity of LGN and its binding partners is imprecise, frequently resulting in oblique orientations at anaphase. In these cases, the apical daughter either retains or loses basement membrane contact following cytokinesis (red or blue nuclei, respectively). If contact is maintained, the apical daughter will reorient into a planar position. In contrast, if contact is lost, the apical daughter further stacks above its basal partner. Upon loss of α-catenin, vinculin, or afadin, telophase reorientation in either direction fails, resulting in persistent oblique divisions. In comparison, LGN loss reduces perpendicular anaphase orientations, and oblique divisions always correct to a final, planar orientation.

### Corrective mechanisms in OCD

Our findings contribute to a growing number of corrective mechanisms which can counterbalance stem cell division orientation errors in order to preserve tissue homeostasis. In *Drosophila* neuroblasts, the “telophase rescue” pathway—mediated by the scaffolding protein Dlg and motor protein Khc73—can compensate for errors in spindle orientation by relocalizing fate determinants, thus preserving normal daughter cell fates (Cai, Chia, & Yang, 2001; Peng, Manning, Albertson, & Doe, 2000; Siegrist & Doe, 2005). However, telophase rescue differs from the telophase correction we report here in that division orientation errors are not corrected in telophase, but rather, the fate determinants themselves are repositioned relative to the new division axis. In the developing epidermis, it has been shown that Insc overexpression can promote apical LGN localization and drive an increase of perpendicular divisions (Poulson & Lechler, 2010; Williams et al., 2011), but that under some circumstances, NuMA can redistribute laterally, perhaps in an effort to prevent the hyper-differentiation that would be driven by excessive asymmetric divisions (Poulson & Lechler, 2010). Our data here, where afadin, α-catenin, or vinculin loss can override the perpendicular-correcting cue provided by LGN, provide a potential molecular explanation for this plasticity.

Other examples of temporally-dynamic OCDs include cyst stem cells of the *Drosophila* testis, which display randomized division orientation until anaphase, at which point one spindle pole becomes anchored at the interface with the niche-defining hub cell, driving division away from the niche (Cheng, Tiyaboonchai, Yamashita, & Hunt, 2011). In addition, dividing cells within the monolayered *Drosophila* follicular epidermis partially extrude during mitosis and frequently demonstrate oblique division angles, which are corrected by reinsertion into the epithelium in an adhesion-dependent manner (Bergstralh, Lovegrove, & St Johnston, 2015). A more extreme example of this extrusion/reinsertion model has been observed in intestinal organoids, where mitotic cells migrate to the luminal surface and undergo planar divisions before reinserting into the epithelium, thereby driving the patterning of secretory and stem cells (McKinley et al., 2018). Furthermore, genetic alterations in MDCK cells—specifically, *LGN* knockdown or *Par1b* overexpression—can drive out-of-plane divisions which are capable of correcting during anaphase via an apical actomyosin compressive force (Lazaro-Dieguez & Musch, 2017; Zheng et al., 2010). Taken together, these studies and ours suggest that many of these corrective mechanisms rely on polarity, cell-adhesion, and actin dynamics. It will be interesting to monitor the continued exploration of these corrective mechanisms and to further probe the cytoskeletal dynamics underlying these phenomena.

## MATERIALS AND METHODS

### Animals

Mice were housed in an AAALAC-accredited (#329; June 2017), USDA registered (55-R-0004), NIH welfare-assured (D16-00256 (A3410-01)) animal facility. All procedures were performed under IACUC-approved animal protocols (16-162). For all live imaging experiments, mT/mG (Gt(ROSA)26Sortm4(ACTB-tdTomato,-EGFP) Luo/J; Jackson Labs #007576 via Liqun Luo, Stanford University) homozygous females with at least one copy of the *K14-Cre* allele (Dassule, Lewis, Bei, Maas, & McMahon, 2000) were crossed to males of the identical genotype. For fixed sample imaging, wild-type CD1 mice (Charles River; #022) were utilized. *Afdn*^*fl*/*fl*^ animals(Beaudoin et al., 2012) were maintained on a mixed C57B6/J CD1 background and either bred to the same *K14-Cre* allele or injected with lentiviral Cre-mRFP1 (see below). The procedure for producing, concentrating and injecting lentivirus into amniotic fluid of E9.5 embryos has been previously described and is briefly detailed below (Beronja et al., 2010).

### Live Imaging

The live imaging protocol used in this study was adapted from the technique recently described by the Devenport lab (Cetera et al., 2018). A 1% agar solution/media solution containing F-media (3:1 DMEM:F12 + 10% FBS + 1% Sodium bicarbonate + 1% Sodium Pyruvate + 1% Pen/Strep/L-glut mix), was cooled and cut into 35mm discs. Epidermal samples measuring ~4-6mm along the AP axis and ~2-3mm along the medial-lateral axis were extracted from the mid-back of E16.5 mT/mG embryos. These explants were placed dermal-side down onto the gel/media disc, then sandwiched between the gas-permeable membrane of a 35mm lumox culture dish (Sardstedt; 94.6077.331). Confocal imaging was performed utilizing a Zeiss LSM 710 Spectral confocal laser scanning microscope equipped with a 40X/1.3 NA Oil Plan Neo objective. Images were acquired with 5 minute intervals and a Z-series with 0.5 µm step-size (total depth ranging from 20-30 microns) for 3-9 hours. Explants were cultured at 37°C with 5.0% CO2 for >1.5 hours prior to- and throughout the course of imaging. Divisions occurring close to the tissue edge or showing any signs of disorganization/damage were avoided to exclude morphological changes associated with wound-repair. 4D image sets were deconvolved using AutoQuant X3 and processed using ImageJ (Fiji).

### Lentiviral Injections

For full protocol, please see Beronja, et al. (ref. 24). This protocol is approved via IACUC #16-162. Pregnant CD1, *mTmG/K14-Cre*, or *Afdn*^*fl*/*fl*^ females were anesthetized and the uterine horn pulled into a PBS filled dish to expose the E9.5 embryos. Embryos and custom glass needles were visualized by ultrasound (Vevo 2100) to guide microinjection of ~0.7 μl of concentrated lentivirus into the amniotic space. Three to ten embryos were injected depending on viability and litter size. Following injection, the uterine horn(s) were reinserted into the mother’s thoracic cavity, which was sutured closed. The incision in the skin was resealed with surgical staples and the mother provided subcutaneous analgesics (5 mg/kg meloxicam and 1-4 mg/kg bupivacaine). Once awake and freely moving, the mother was returned to its housing facility for 5-7 days, at which point E14.5-16.5 embryos were harvested and processed accordingly.

### Constructs and RNAi

For afadin and vinculin RNAi targeting, we tested ~10 shRNAs for knockdown efficiency in primary keratinocytes. These sequences were selected from The RNAi Consortium (TRC) Mission shRNA library (Sigma) versions 1.0, 1.5, and 2.0 and cloned using complementary annealed oligonucleotides with AgeI/EcoRI linkers. For LGN and α-catenin, we utilized an shRNA that had been previously validated with our lentiviral injection technique(Beronja et al., 2010; Williams et al., 2011). shRNA clones are identified by the gene name with the nucleotide base (NCBI Accession number) where the 21-nucleotide target sequence begins in superscript (e.g. *Afdn*^*2711*^). Lentivirus was packaged in 293FT or TN cells using the pMD2.G and psPAX2 helper plasmids (Addgene plasmids #12259 and #12260, respectively). For knockdown screening, primary keratinocytes were seeded at a density of ~150,000 cells per well into 6-well plates and grown to ~80% confluency in E-Low calcium medium and infected with an MOI of ~1. Approximately 48 h post-infection, keratinocytes were treated with puromycin (2 μg/mL) to generate stable cell lines. After 3-4 days of puromycin selection, cells were lysed and RNA isolated using the RNeasy Mini Kit (Qiagen). cDNA was generated and amplified from 10-200 μg total RNA using either Superscript VILO (Invitrogen) or iScript (Bio-Rad). mRNA knockdown was determined by RT-qPCR (Applied Biosystems 7500 Fast RT-PCR) using 2 independent primer sets for each transcript with *Hprt1* and cyclophilin B (*Ppib2*) as reference genes and cDNA from stable cell lines expressing Scramble shRNA as a reference control. Primer efficiencies were determined using dose-response curves and required to be >1.8, with relative transcript abundance determined by the ΔΔCT method. RT-qPCR runs were performed in triplicate with the mean knockdown efficiency determined by calculating the geometric mean of the ΔΔCT values for at least two independent technical replicates. The following primer sequences were used: *Afdn* (fwd-1: 5’-ACGCCATTCCTGC-CAAGAAG -3’, rev-1: 5’-GCAAAGTCTGCGGTATCGGTAGTA -3’; fwd-2: 5’-GGGGATGACAGGCTGATGAAA -3’, rev-2: 5’-CGATGCCGCT-CAAGTTGGTA -3’), *Vcl* (fwd-1: 5’-TACCAAGCGGGCACTTATTCAGT-3’, rev-1: 5’-TTGGTCCGGCCCAGCATA -3’; fwd-2: 5’-AAGGCTGTG-GCTGGAAACATCT -3’, rev-2: 5’-GGCGGCCATCATCATTGG -3’). The following shRNA targeting sequences were used: *Afdn*^*2711*^ (5’-CCTGAT-GACATTCCAAATATA -3’), *Vcl*^*3466*^ (5’-CCCTGTACTTTCAGTTACTAT-3’), *Vcl*^*2803*^ (5’-CCACGATGAAGCTCGGAAATG -3’), *Ctnna1*^*912*^ (5’-CGCTCTCAACAACTTTGATAA -3’), *LGN*^*1617*^ (5’-GCCGAATTGGAA-CAGTGAAAT -3’), *Scramble* (5’-CAACAAGATGAAGAGCACCAA-3’).

### Antibodies, immunohistochemistry, and fixed imaging

E14.5 embryos were mounted whole in OCT (Tissue Tek) and frozen fresh at −20°C. E16.5 embryos were skinned and flat-mounted on Whatman paper. In both cases, infected and uninfected littermate controls were mounted in the same blocks to allow for direct comparisons on the same slide. For α-tubulin staining of metaphase spindles, samples were kept warm and pre-fixed with room-temperature 4% paraformaldehyde for 10 minutes before OCT embedding. Frozen samples were sectioned (8 μm thick) on a Leica CM1950 cryostat, mounted on SuperFrost Plus slides (ThermoFisher) and stored at −80°C. For staining, sections were thawed at 37°C for 5-15 min, fixed for 5 min with 4% paraformaldehyde, washed with PBS and blocked for 1h with gelatin block (5% NDS, 3% BSA, 8% cold-water fish gelatin, 0.05% Triton X-100 in PBS). Primary antibodies were diluted in gelatin block and incubated overnight in a humidity chamber at 4°C. Slides were then washed with PBS and incubated with secondary antibodies diluted in gelatin block at room temperature (~25°C) for 2 hours, counterstained with DAPI (1:2000) for 5 minutes and mounted in ProLong Gold (Invitrogen). Actin was visualized by phalloidin-AF647 staining (Life Technologies; 1:500) simultaneously with secondary antibody incubation. Images were acquired using LAS AF software on a Leica TCS SPE-II 4 laser confocal system on a DM5500 microscope with ACS Apochromat 20x/0.60 multi-immersion, ACS Apochromat 40x/1.15 oil, or ACS Apochromat 63x/1.30 oil objectives.

The following primary antibodies were used: survivin (rabbit, Cell Signaling 2808S, 1:500), LGN(Williams et al., 2011) (guinea pig, 1:500), LGN (rabbit, Millipore ABT174, 1:2000), phospho-histone H3 (rat, Abcam ab10543, 1:1,000), mCherry (rat, Life Technologies M11217, 1:1000-3000), α4-integrin (rat, ThermoFisher 553745, 1:1,000), Gαi3 (rabbit, EMD Millipore 371726, 1:500), GFP (chicken, Abcam ab13970, 1:1,000), dynactin (goat, Abcam ab11806, 1:500), NuMA (mouse IgM, BD Transduction Labs 610562, 1:300), α-tubulin (rat, EMD Millipore CBL270, 1:500), pericentrin (rabbit, Covance PRB-432C, 1:500), Par3 (rabbit, EMD Millipore 07-330, 1:500), E-cadherin (rat, Life Technologies 131900, 1:1,000), E-cadherin (goat, R&D System AF748, 1:1,000), α-catenin (rabbit, Invitrogen 71-1200, 1:300) α18 (rat, generous gift of Dr. Nagafuchi at Nara Medical University, 1:10,000), vinculin (mouse IgG, Sigma V9131, 1:500), vinculin (rabbit, generous gift of Dr. Keith Burridge at University of North Carolina, 1:1000), afadin (rabbit, Sigma A0224, 1:500).

The following secondary antibodies were used (all antibodies produced in donkey): anti-rabbit AlexaFluor 488 (Life Technologies, 1:1000), anti-rabbit Rhodamine Red-X (Jackson Labs, 1:500), anti-rabbit Cy5 (Jackson Labs, 1:400), anti-rat AlexaFluor 488 (Life Technologies, 1:1000), anti-rat Rhodamine Red-X (Jackson Labs, 1:500), anti-rat Cy5 (Jackson Labs, 1:400), anti-guinea pig AlexaFluor 488 (Life Technologies, 1:1000), anti-guinea pig Rhodamine Red-X (Jackson Labs, 1:500), anti-guinea pig Cy5 (Jackson Labs, 1:400), anti-goat AlexaFluor 488 (Life Technologies, 1:1000), anti-goat Cy5 (Jackson Labs, 1:400), anti-mouse IgG AlexaFluor 488 (Life Technologies, 1:1000), anti-mouse IgG Cy5 (Jackson Labs, 1:400), anti-mouse IgM Cy3 (Jackson Labs, 1:500).

### Keratinocyte culture and Calcium-shift assays

Primary mouse keratinocytes were maintained in E medium with 15% chelated FBS and 50 μM CaCl (E low medium). For viral infection, keratinocytes were plated at ~150,000 cells per well in a 6-well plate and incubated with lentivirus in the presence of polybrene (1 μg/mL) and centrifuged at 1,100 xg for 30 min at 37°C. Stable cell lines were generated/maintained by adding puromycin (2 μg/mL) 48 h after infection and continual antibiotic treatment following. Calcium shifts were performed by seeding ~45,000 cells per well into 8-well Permanox chamber slides (Lab-Tek 177445) coated with poly-L-lysine, collagen, and fibronectin. Once cells reached ~85% confluency (~12-16 hours) cells were switch to high Ca^2+^ (1.5mM) medium and grown for the indicated period of time (30 min to 8 hours). Cells were fixed with 4% paraformaldehyde in PBS warmed to room-temperature. Immunostaining was performed using the same protocol as for slides (see above).

### Measurements, quantification, graphing, and statistics

All image analysis was performed in Fiji (ImageJ). Mitotic cells in metaphase were identified based on nuclear morphology. Metaphase spindle orientation was measured as the angle between a vector orthogonal to the metaphase plate and parallel to the basement membrane. Anaphase cells were identified by both nuclear condensation and widely distributed surviving staining between daughter cells. Telophase cells were distinguished due to reduced nuclear condensation and dual-punctate survivin staining. Division orientation was measured as the angle between a vector connecting the center of each daughter nucleus and a vector running parallel to the basement membrane. The same methodology was used to measure division orientation in live imaging experiments. In cases lacking nuclear labeling, the position of the nuclei was inferred based on cell volume/shape changes as demonstrated in Fig. 1e and Supplementary Fig. 1. Telophase correction (ϕ-θ) was quantified as the difference between division orientation at anaphase onset (ϕ) and division orientation 1 h later (θ). The presence of basal contact for the more apical daughter was determined by analyzing cell morphology in both en-face and orthogonal perspectives as in Fig. 1e.

Quantification of fluorescence intensity in adhesion assays was performed by orthogonal linescans at three positions along the junction length (~25th, 50th, and 75th quarter). In cases where junctions appeared punctate, discreet puncta were evaluated to avoid measuring regions lacking junction formation. Signal centers were set based on maximum intensity of either E-cadherin or α-catenin (where appropriate). To quantify ratios, the geometric mean fluorescent intensities of the 3 values nearest the junction center were used. Quantification of junction continuity was performed by linescans of E-cadherin fluorescence intensity along the entire length of the junction, excluding the vertex of multiple cells (i.e. tricellular junctions). We then calculated % of these intensity measurements above a threshold, which was evaluated for each individual junction using the mean center intensity of three orthogonal scans described earlier in this paragraph.

LGN localization patters (e.g. apical, weak/absent, or other) were determined for cells labeled with pHH3, irrespective of the lentiviral H2B-RFP reporter to avoid bias. Imaging was performed with WT controls and experimental samples on the same slide to avoid variation in antibody staining. Radial localization of LGN was measured by determining the angle between two vectors: one drawn from the LGN signal center to the center of the nucleus, the other drawn parallel to the basement membrane. Crescents oriented at the apical side were given positive values, while those at the basal side were given negative values. Radial variance between LGN signal and spindle or division axis were determined by drawing two vectors: one for the radial orientation of the LGN signal center and a second between either the spindle poles or between the center of the daughter nuclei (as demonstrated in Fig 6d,f). Radial fluorescent intensity values were measured by linescans originating at the site of basement membrane contact and tracing the edge of the cell. Each measurement along the length of the scan was then set as a part of whole, operating with the assumption that ~50% of the total length would represent the apical surface. Data displayed in Supplementary Fig. 6b, g are mean with 95% confidence interval.

Local cell density was calculated by counting the number of neighboring cells and dividing this number by their area. Area was measured in a single z-plane determined to be the cell center by orthogonal slices. Regions where the tissue sloped at an extreme angle were excluded due to inability to capture cell centers for all neighbors.

All statistical analyses and graphs were generated using GraphPad Prism 8 and Origin 2015 (OriginLab). Error bars represent standard error of the mean (s.e.m.) unless otherwise noted. Statistical tests of significance were determined by Mann-Whitney U-test (non-parametric) or student’s t-test (parametric) depending on whether the data fit a standard distribution (determined by pass/fail of majority of the following: Anderson-Darling, D’Agostino & Pearson, Shapiro-Wilk, and Kolmogorov-Smirnov tests). Cumulative frequency distributions were evaluated for significant differences by Kolmogorov-Smirnov test. χ^2^ tests were utilized to evaluate expected (control) against experimental distributions of categorical values (e.g. LGN apical/absent/other distributions in Fig. 4f). All box-and-whisker plots are displayed as Tukey plots where the box represents the interquartile range (IQR, 25th-75th percentiles) and the horizontal line represents the median. Whiskers represent 1.5xIQR unless this is greater than the min or max value. Figures were assembled using Adobe Photoshop and Illustrator CC 2017.

## Acknowledgements

We thank members of the Williams and Peifer Labs for their critical feedback throughout the process. We thank Dr. Danelle Devenport (Princeton) for graciously sharing her live imaging protocol. We thank Dr. Akira Nagafuchi (Nara Medical University) for sharing the α18 antibody. We thank Dr. Brent Hoffman and Evan Gates (Duke University) for their effort and feedback regarding E-cadherin mediated junctional tension. We thank Dr. Keith Burridge (UNC) for sharing the vinculin (Rb) polyclonal antibody. We thank Kendra Niederkorn for design input into the model presented in Fig. 8. KJL was supported by an NIH Ruth L. Kirschstein Predoctoral National Research Service Award (F31 DE026956). KMB was supported by an NIH/NIDCR K08 Mentored Clinical Scientist Research Career Development Award (DE026537). SW was supported by a Sidney Kimmel Scholar Award (SKF-15-165).

## Author Contributions

KJL and SEW designed, and KJL, CPD, DCS, and AJB performed the experiments. KJL and SEW analyzed data. LFK provided Afdnfl/fl mice for afadin knockout experiments. KMB provided extensive intellectual input at project onset and throughout the course of the work. KJL and SEW made the figures and wrote the manuscript with input from all authors.

---

**Figure S1.**
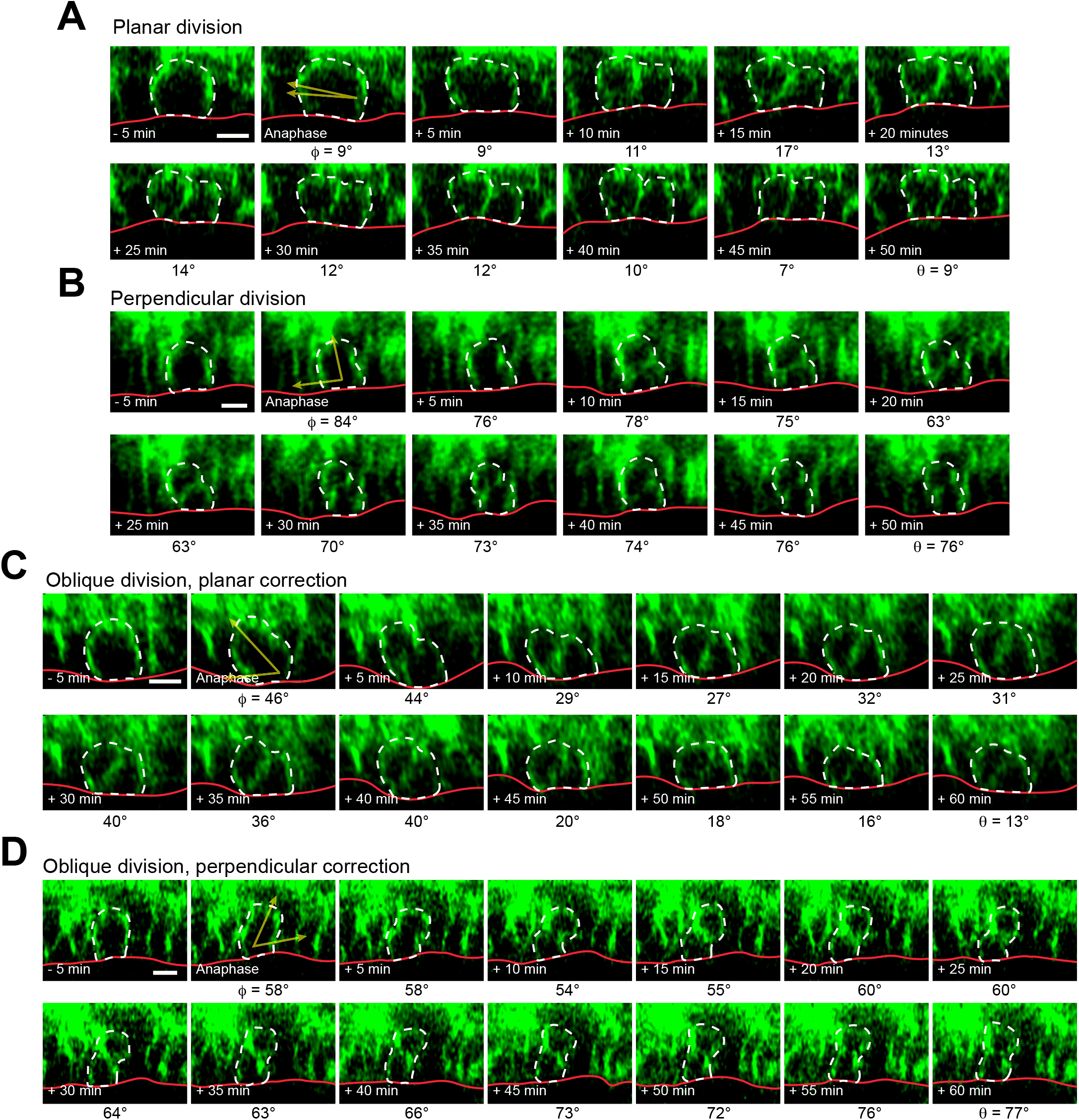
Related to Figure 1 | Telophase reorientation behaviors in E16.5 wild-type epidermal explants. Stills (z-projections from 0.5 µm steps) from movies of mitotic cells, acquired at 5 minute intervals. Image series depict a division that initiates and completes in a planar orientation (**A**); initiates and completes in a perpendicular orientation (**B**); or initiates at an oblique orientation and corrects to planar (**C**) or perpendicular (**D**). Division angles (yellow arrows in images at anaphase onset, ϕ), relative to epidermal-dermal junction (red) are shown below each still. Scale bars, 10μm.

**Figure S2.**
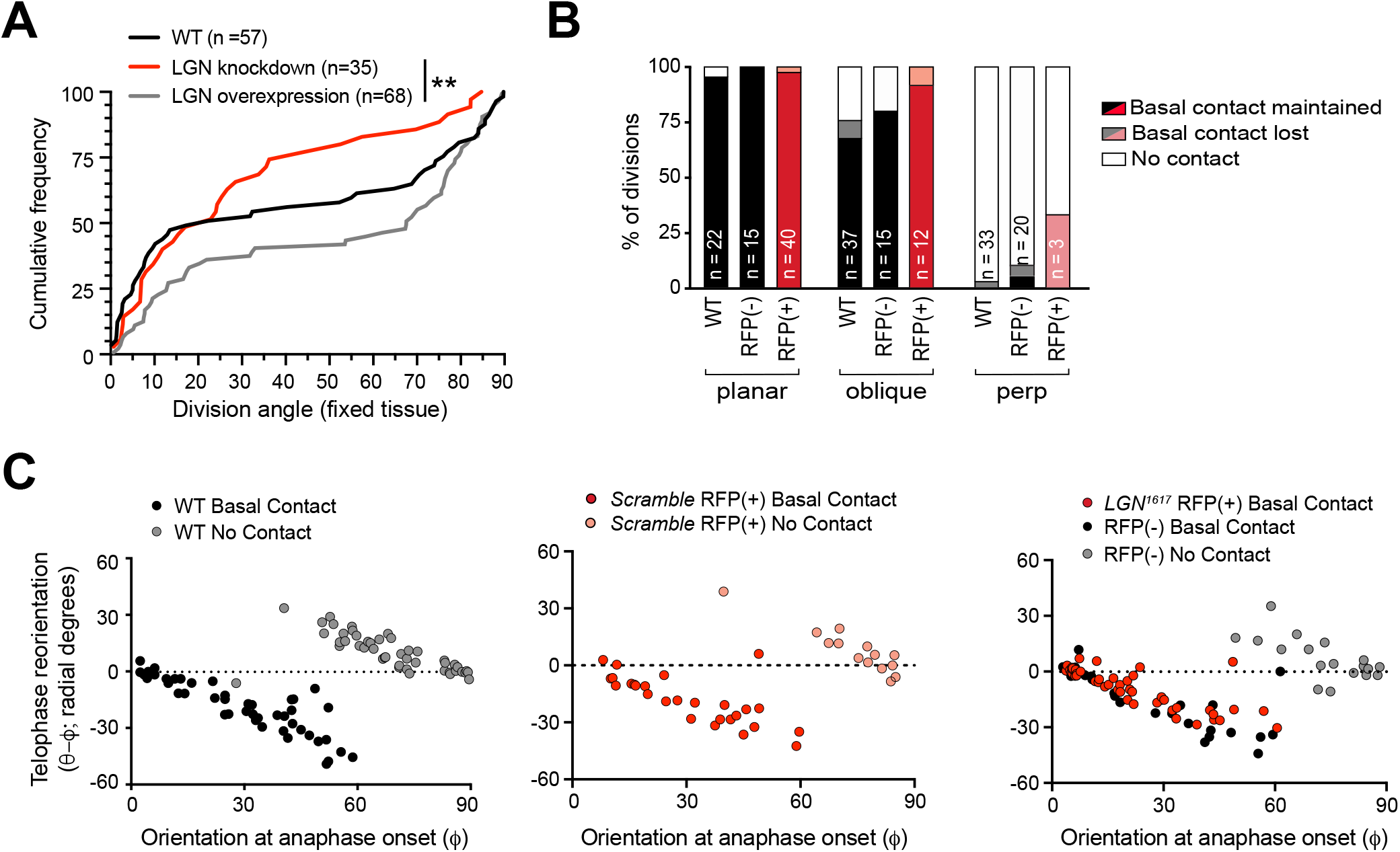
Related to Figure 2 | Scramble controls display normal telophase reorientation while LGN knockdown results in only planar-directed reorientation. (**A**) Cumulative frequency distribution of telophase division orientation from wild-type littermates (black), *LGN* knockdown (*LGN*^*1617*^ RFP+, red), and *LGN* overexpression (*LGN*^*1617*^ RFP-*LGN*^*1617*^, grey) cells from E16.5 fixed sagittal sections. *LGN* loss biases distribution toward planar divisions, while *LGN* overexpression biases toward perpendicular divisions; n indicates divisions measured from 3-6 independent embryos. (**B**) On the x-axis, divisions are grouped by orientation at anaphase onset (ϕ, planar = 0-30°; oblique = 30-60°; perpendicular = 60-90°) and *LGN*^*1617*^ status. The y-axis depicts the proportion of divisions that lack (white), maintain (black/red), or lose (grey/pink) basal contact in the following hour. ~95% of planar divisions initiate and maintain basal contact; oblique divisions make and initiate contact less frequently (~75% of the time); while perpendicular divisions almost never make contact. *LGN* knockdown (red bars) does not alter this behavior. (**C**) Radial anaphase correction (ϕ - θ) plotted versus initial anaphase orientation. Cells where apical daughter basal contacts were detected are shown as black/red circles, while those lacking basal contacts are shown in grey/pink. In wild-type controls (left) basal contact correlates with planar reorientation, while lack of contact results in perpendicular reorientation. Telophase correction (ϕ - θ) in *Scramble* RFP+ cells (middle panel), grouped by basal contact status. Like wild-type cells, *Scramble* RFP+ cells undergo telophase reorientation in a basal contact-dependent manner. Both *LGN*^*1617*^ RFP+ (right panel; red) and wild-type RFP-negative cells (black) with basal contacts correct to planar, demonstrating that LGN is not required for this behavior. *LGN* knockdown cells only rarely (n=3) lack basement membrane contact, so this group is not shown. Scale bars, 10μm. * P < 0.05, ** P < 0.01 by Kolmogorov-Smirnov test.

**Figure S3.**
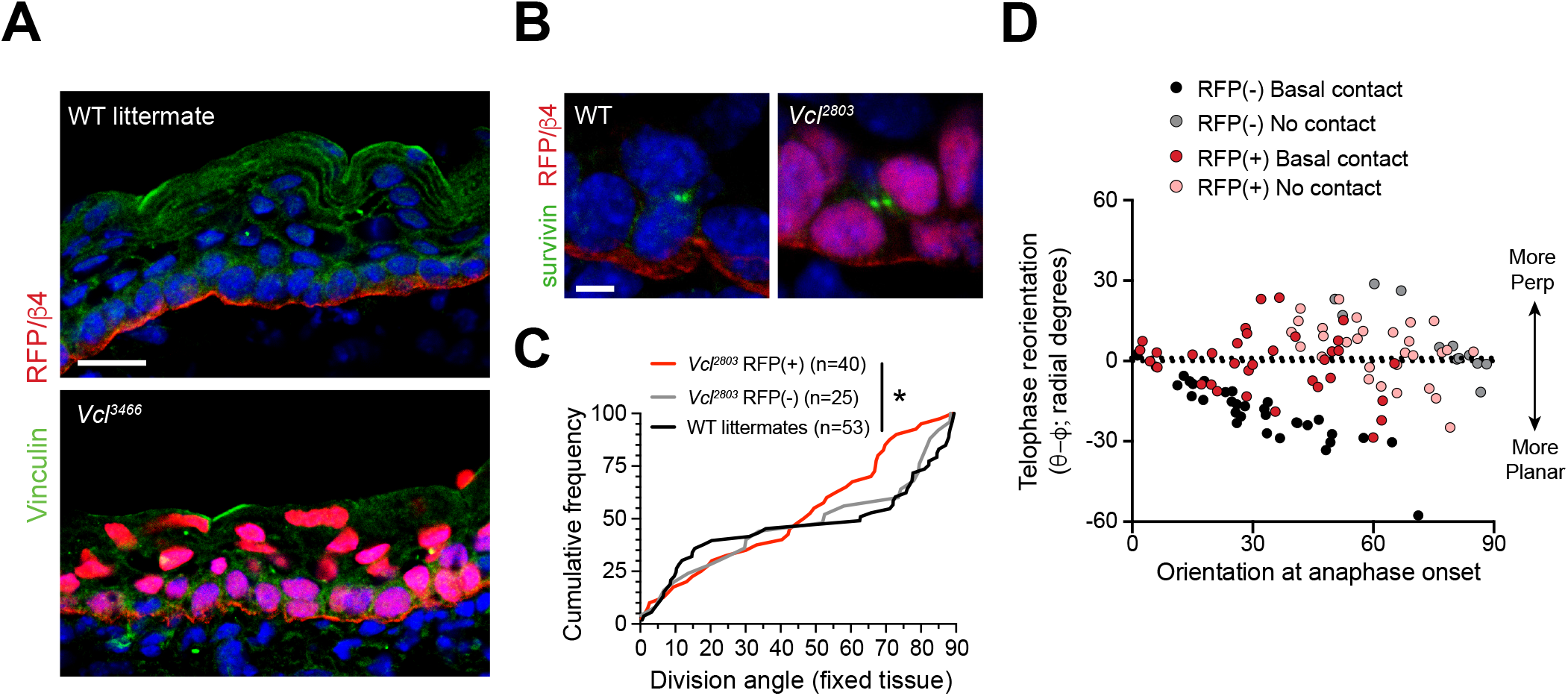
Related to Figure 3 | Validation of Vcl shRNAs. (**A**) E16.5 epidermis infected with *Vcl*^*3466*^ H2B-RFP (red) and stained rabbit with anti-vinculin antibody. While suprabasal staining is dramatically reduced in infected samples, some non-specific basal-layer staining remains. (**B**) Telophase cells marked by survivin (green), from E16.5 wild-type littermates (left) and *Vcl*^*2803*^ RFP+ knockdowns (right). (**C**) Cumulative frequency distribution of division orientation in E16.5 *Vcl*^*2803*^ H2B-RFP mosaic epidermis. Compare to *Vcl*^*3466*^ shRNA, shown if Fig. 3b. (**D**) Radial telophase correction (ϕ - θ) plotted versus initial anaphase orientation (ϕ), for *Vcl*^*3466*^ RFP+ (red/pink) and RFP-negative (black/grey) cells, binned based on presence (black/red) or absence (grey/pink) of basal contact maintenance. Vinculin knockdown cells are unresponsive to basal contact and show no obvious correction pattern. Scale bar, 20μm (**A**), 5μm (B). * P < 0.05 by Kolmogorov-Smirnov test.

**Figure S4.**
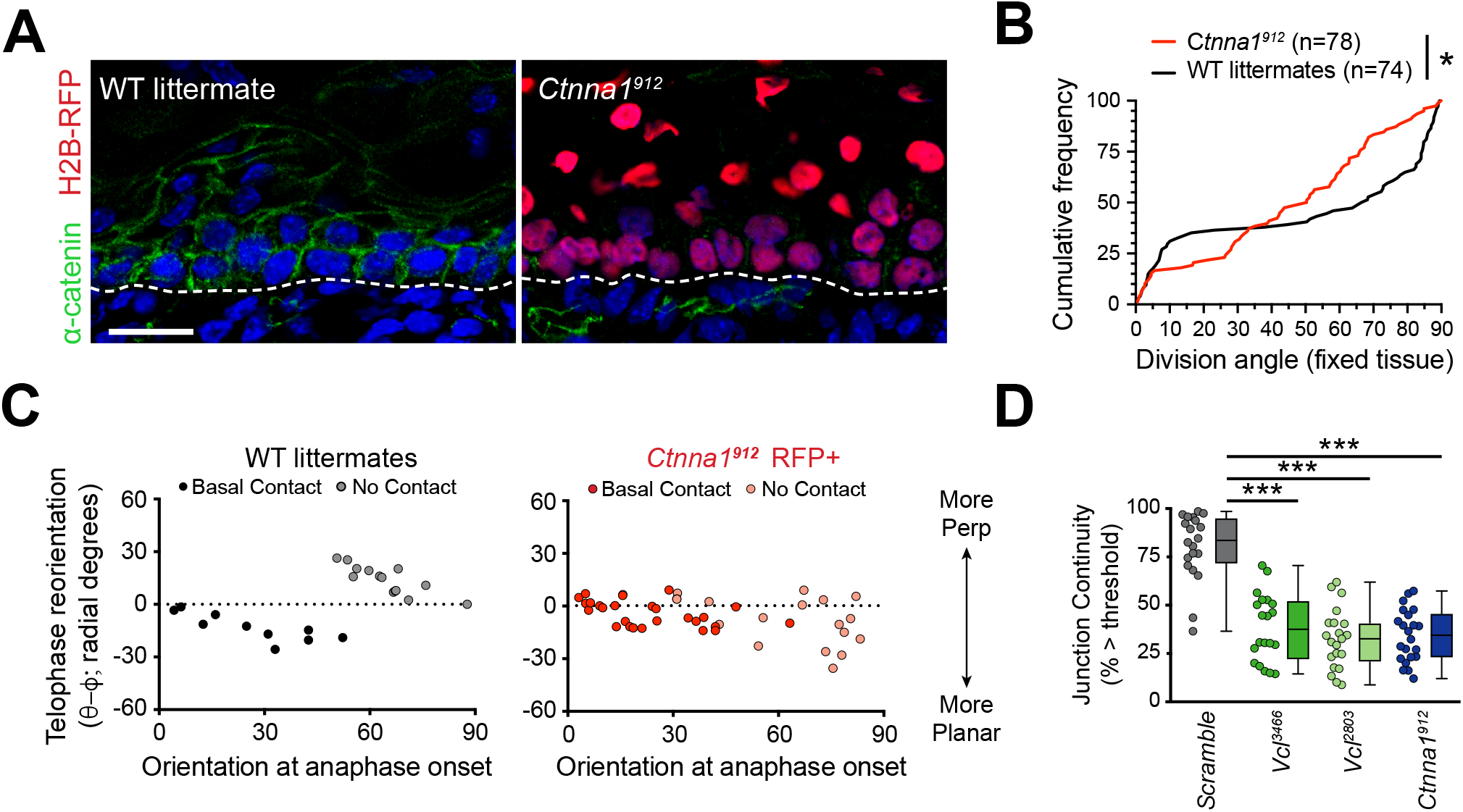
Related to Figure 4 | α-catenin knockdown is effective *in vivo* and results in basal-contact independent reorientation. (**A**) Immunofluorescent images taken from E16.5 sagittal sections of wild-type littermate controls (left) or transduced with *Ctnna1*^*912*^ H2B-RFP (right). Epidermal junctional α-catenin (green) is lost in *Ctnna1*^*912*^ RFP+ epidermis. (**B**) Cumulative frequency distributions of telophase division angles from E16.5 sections of *Ctnna1*^*912*^ knockdown (red) and control littermates (black); n indicates measurements from 6-7 independent embryos. (**C**) Radial change in division angle from anaphase onset to one hour later (ϕ - θ, y-axis), plotted relative to initial anaphase orientation (ϕ, x-axis) for wild-type littermates (left) and *Ctnna1*^*912*^ knockdown embryos (right). Knockdown of α-catenin results in minimal correction, particularly for cells that maintain basal contact. (**D**) Quantification of junction continuity, based on fluorescent intensity of E-cad puncta, after an 8h Ca^2+^ shift. Knockdown of *Vcl* or *Ctnna1* results in punctate junction morphology, a characteristic of immature spot junctions (see Supplementary Fig. 7A-C).

**Figure S5.**
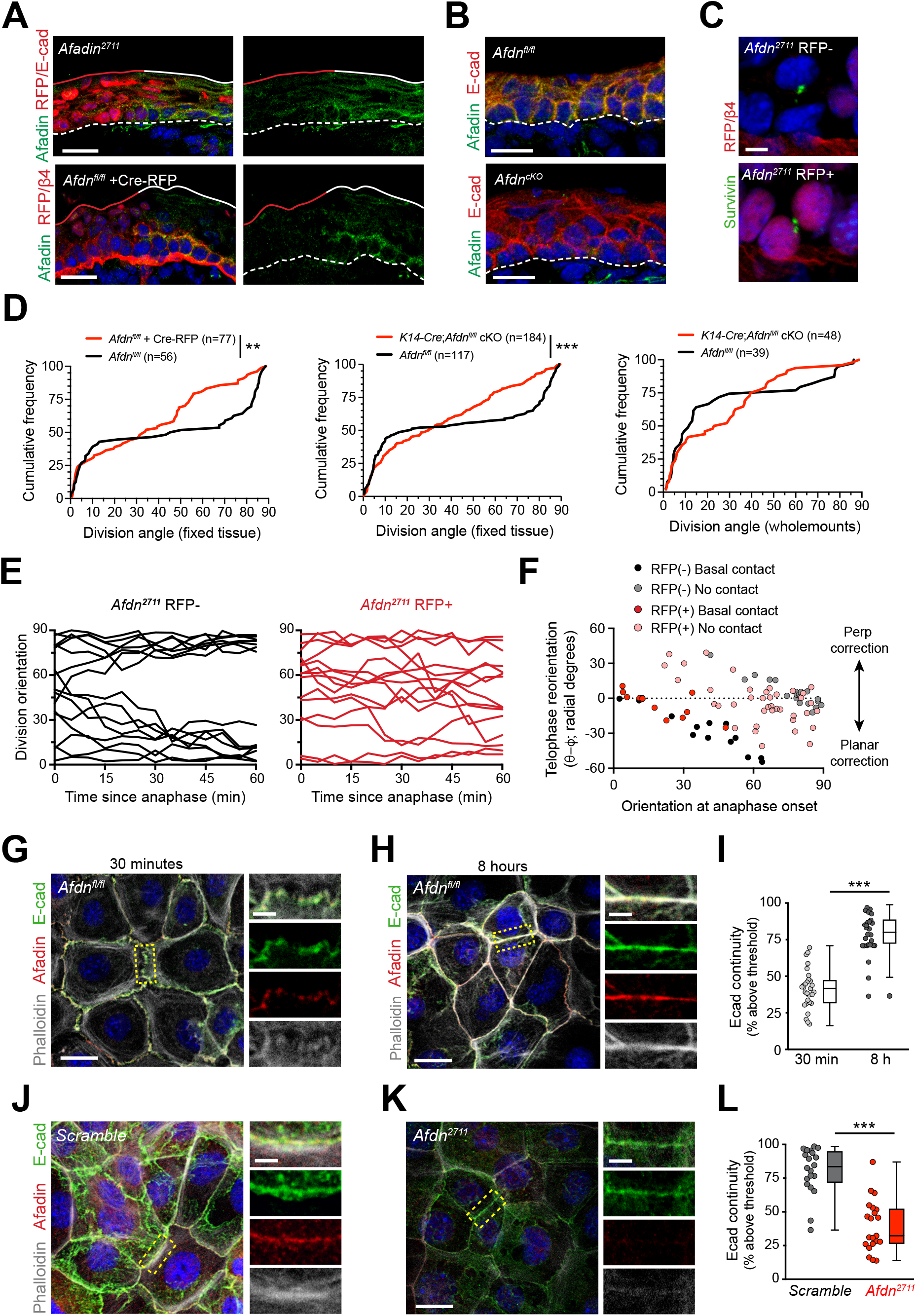
Related to Figure 5 | Multiple models of afadin loss-of-function demonstrate telophase division orientation defects. (**A-B**) Afadin (green) and E-cadherin (red) immunostaining in E16.5 sections. (**A**) Mosaic region of *Afdn*^*2711*^ H2B-RFP (top panel) or Cre-RFP (in *Afdn*^*fl*/*fl*^ embryo; bottom panel) lentiviral transduction. Region of high transduction (red line) demonstrates efficient loss of junctional afadin signal, spared in region of low transduction (white line). (**B**) E16.5 *Afdn*^*fl*/*fl*^ controls (left) with conditional deletion mediated by *K14-Cre* (*cKO*). (**C**) RFP-negative (top) and RFP+ (bottom) telophase cells from mosaic *Afdn*^*2711*^ epidermis, identified by survivin expression (green). (**D**) Cumulative frequency distributions of telophase division angles from E16.5 sagittal sections, or wholemounts (far right panel). *Afdn* knock-down or knockout results in randomized division orientation at telophase, regardless of Cre driver used or analysis method; n indicates number of observed divisions from 3-4 independent embryos. Note, fewer perpendicular divisions are observed in wholemounts compared to sections, likely due to 1) the relative difficulty of detecting perpendicular divisions compared to planar divisions in wholemounts, and 2) the likelihood of undercounting planar-mediolateral divisions in sagittal sections. (**E**) Timelines of division orientation at 5-minute intervals from movies of *Afdn*^*2711*^ RFP-negative (left) and RFP+ (right) for 15 representative cells per group. Telophase reorientation establishes bimodal distribution within ~30 minutes in RFP-control cells that enter anaphase at oblique angles, while RFP+ cells fail to demonstrate any sorting behavior over a full hour following anaphase onset. (**F**) Radial correction (ϕ - θ) for *Afdn*^*2711*^ RFP-negative (black/grey) and RFP+ (red/pink) plotted versus initial anaphase orientation (ϕ). *Afadin* knockdown cells are less responsive to the orienting function of basal contacts compared to RFP-negative controls. (**F**) Basal contact maintenance over a 1 h interval post-anaphase for *Afdn*^*2711*^ RFP-negative (black) and RFP+ (red) cells. *Afdn* knockdown apical daughter cells, particularly those that enter anaphase at oblique angles, fail to maintain basal contact through mitosis. (**G-H**) Timecourse of junction formation in wild-type *Afdn*^*fl*/*fl*^ primary keratinocytes 30 min (**G**) and 8 h (**H**) after the addition of 1.5 mM Ca2+, stained for E-cad (green), afadin (red), and phalloidin (grey). Between early and late timepoints, junctions transform from punctate to linear and actin becomes tightly associated with the junction. (**I**) Quantification of E-cad continuity along junction length in *Afdn*^*fl*/*fl*^ primary keratinocytes at 30 min and 8 h timepoints. (**J-K**) *Scramble* (**J**) and Afadin knockdown (*Afdn*^*2711*^; **K**) keratinocytes after 8 h Ca^2+^ shift, stained same as in (**G-H**). (**L**) Quantification of E-cad continuity in *Scramble* and *Afdn*^*2711*^ as in (**I**). Scale bars, 20µm (**A-B,G-H,J-K**), 5µm (**C**). ** P < 0.01, *** P < 0.001, determined by Kolmogorov-Smirnov test (**D**) or student’s t-test (**I,L**).

**Figure S6.**
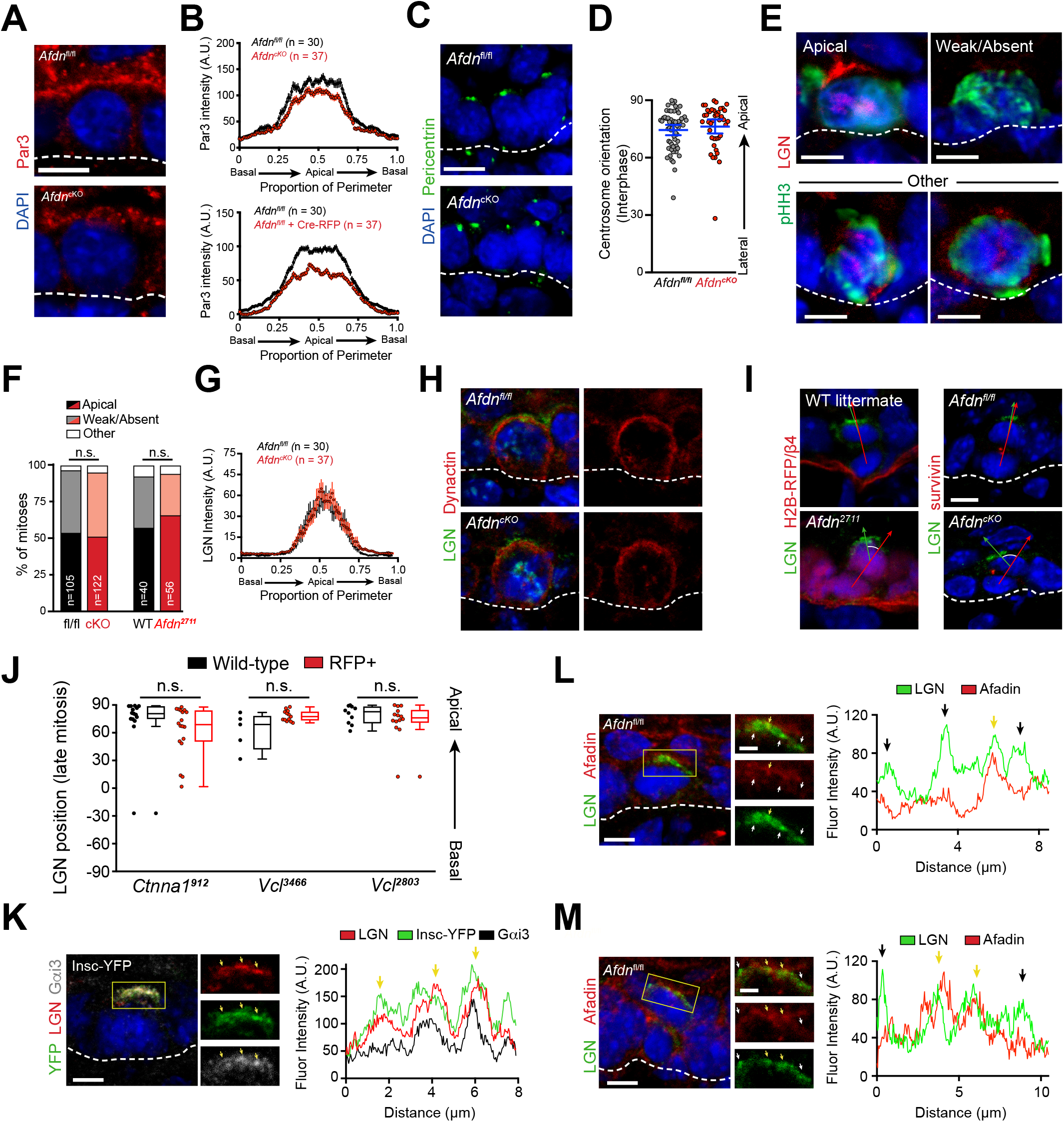
Related to Figure 6 | Afadin loss-of-function division orientation phenotypes are polarity and LGN-independent. (**A**) Immunostaining of E16.5 interphase cell with Par3 (red). (**B**) Quantification of Par3 radial fluorescent intensity. Apical accumulation is reduced by *afadin* knockout via *K14-Cre* (*Afdn*^*cKO*^) or lentiviral Cre-RFP; n indicates interphase cells from 2-3 independent embryos. (**C**) E16.5 sagittal sections show that centrosomes (green) localize to the apical cortex of interphase cells in both *Afdn*^*cKO*^ basal progenitors and *Afdn*^*fl*/*fl*^ controls. (**D**) Quantification of centrosome radial position in basal keratinocytes; n indicates interphase cells from 2 independent embryos. (**E**) LGN (red) localization patterns in mitotic (green) basal keratinocytes. (**F**) Quantification of LGN rate of recruitment, binned by genotype. LGN localizes to the apical cortex in ~50% of mitoses (black/red), is absent in ~45% (grey/pink), and “other” in the remaining ~5% (white); n indicates mitotic cells from 2-3 independent embryos. (**G**) Quantification of LGN radial fluorescence intensity in E16.5 mitotic cells (as in **B**); n indicates LGN^+^ mitoses from 2-3 independent embryos. (**H**) Dynactin (red) localization in LGN^+^ mitoses is unaffected by afadin knockout. (**I**) *Afdn* knockdown increases radial deviation between LGN (green) and the division plane (red line) (**J**) Quantification of LGN radial localization in telophase (survivin+) mitoses. LGN apical bias is unaffected by α-catenin or vinculin knockdown. (**K**) Costaining for LGN (red), Gαi3 (grey), and mInsc-YFP (green) in E17.5 CD1 embryos infected with mInsc-YFP demonstrates a high degree of colocalization (yellow arrows). Fluorescence intensity linescan across LGN crescent is displayed in the right panel. (**L-M**) Costaining of LGN (green) and afadin (red) in metaphase (**L**) and telophase (**M**) mitoses from E16.5 *Afdn*^*fl*/*fl*^ epidermis. Fluorescence intensity linescan across LGN crescent is displayed in the right panel. Regions of colocalization (yellow arrows) are just as prevalent as regions lacking signal overlap (black arrows). Scale bars, 5μm. P values determined by student’s t-test or Mann-Whitney test depending on tests of normality (**D,J**) or χ^2^ (**F**). n.s. not significant (P > 0.05).

**Figure S7.**
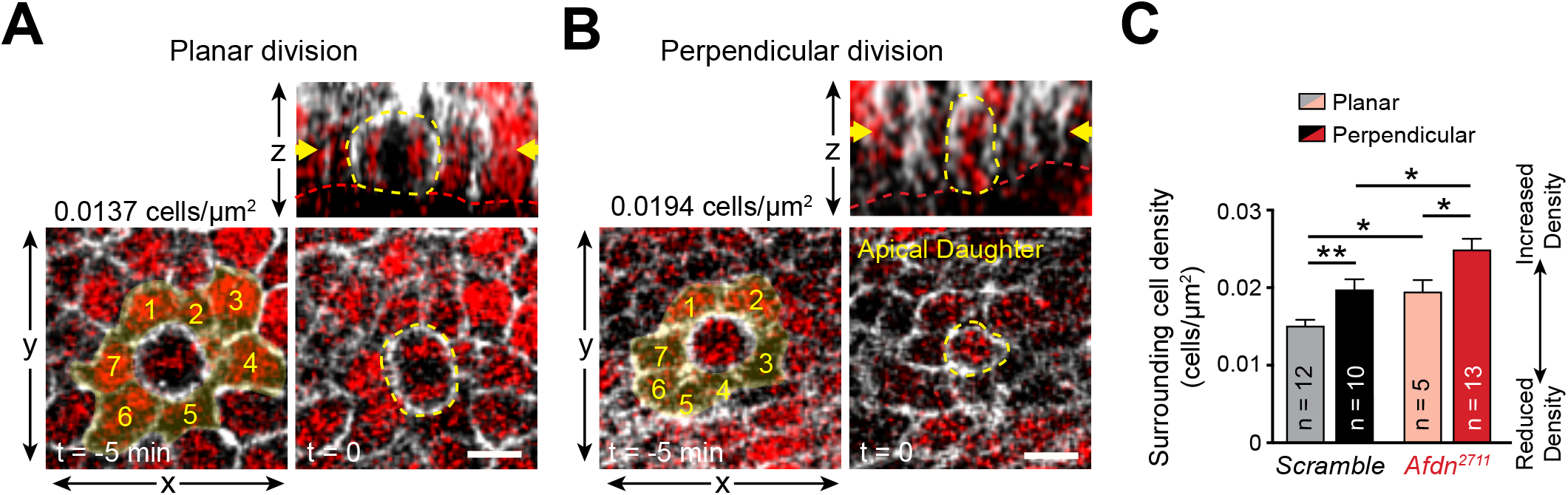
Related to Figure 7 | Afadin knockdown does not perturb density-dependent division orientation behavior. (**A,B**) Relationship between mitotic neighbor cell density and division orientation. (**A-B**) En-face (bottom) and z-projection (top) stills from movies of *Scramble* control epidermis just before (t = −5 min) and at anaphase onset (t = 0). (**C**) Plot of cell density surrounding mitotic events (e.g., the area of cells 1-7 in (**A**) at t=−5 min; highlighted in yellow), grouped by eventual division orientation and genotype. Planar anaphase orientations occur in less dense regions for both *Scramble* and *Afdn*^*2711*^ cells; n indicates divisions observed from 2-4 independent embryos. Scale bars, 10μm. P values determined by student’s unpaired t-test. * P < 0.05, ** P < 0.01.

## SUPPLEMENTAL VIDEO LEGENDS

**Supplemental Video 1. Related to Figure 1 | Planar anaphase orientation is fixed**

Z-projection of *ROSA mTmG K14-Cre* epidermal explants taken from E16.5 embryos imaged in 5 minute intervals. The mitotic cell (white dashed outline) orients its division parallel to the basement membrane (red) resultig in a persistent planar OCD.

**Supplemental Video 2. Related to Figure 1 | Perpendicular anaphase orientation is fixed**

Z-projection of *ROSA mTmG K14-Cre* epidermal explants taken from E16.5 embryos imaged in 5 minute intervals. The mitotic cell (white dashed outline) orients its division orthogonal to the basement membrane (red) resultig in a persistent perpendicular OCD.

**Supplemental Video 3. Related to Figure 1 | Oblique anaphase orientations undergo planar telophase reorientation**

Z-projection of *ROSA mTmG K14-Cre* epidermal explants taken from E16.5 embryos imaged in 5 minute intervals. The mitotic cell (white dashed outline) initiates anaphase at an oblique (30-60°) orientation relative to the basement membrane (red). However, the apical daughter retains basement membrane contact (yellow arrowhead) and corrects into a planar orientation within one hour.

**Supplemental Video 4. Related to Figure 1 | Oblique anaphase divisions display perpendicular correction**

Z-projection of *ROSA mTmG K14-Cre* epidermal explants taken from E16.5 embryos imaged in 5 minute intervals. The mitotic cell (white dashed outline) initiates anaphase at an oblique (30-60°) orientation relative to the basement membrane (red). However, the apical daughter corrects into a perpendicular orientation within one hour.

**Supplemental Video 5. Related to Figure 5 | Oblique anaphase divisions are not corrected in Afdn2711 knockdown**

Z-projection of *ROSA mTmG K14-Cre* epidermal explants taken from E16.5 embryos infected with *Afdn*^*2711*^ H2B-RFP and imaged in 5 minute intervals. The mitotic cell initiates anaphase at an oblique (30-60°) orientation relative to the basement membrane (white dashed line). The apical retains basemement membrane contact (white arrow), but fails to maintain this over time.

## REFERENCES

Beaudoin, G. M., 3rd, Schofield, C. M., Nuwal, T., Zang, K., Ullian, E. M., Huang, B., & Reichardt, L. F. (2012). Afadin, a Ras/Rap effector that controls cadherin function, promotes spine and excitatory synapse density in the hippocampus. J Neurosci, 32(1), 99–110. doi:10.1523/JNEUROSCI.4565-11.2012

Benham-Pyle, B. W., Pruitt, B. L., & Nelson, W. J. (2015). Cell adhesion. Mechanical strain induces E-cadherin-dependent Yap1 and beta-catenin activation to drive cell cycle entry. Science, 348(6238), 1024–1027. doi:10.1126/science.aaa4559

Bergstralh, D. T., Lovegrove, H. E., & St Johnston, D. (2015). Lateral adhesion drives reintegration of misplaced cells into epithelial monolayers. Nat Cell Biol, 17(11), 1497–1503. doi:10.1038/ncb3248

Beronja, S., Livshits, G., Williams, S., & Fuchs, E. (2010). Rapid functional dissection of genetic networks via tissue-specific transduction and RNAi in mouse embryos. Nat Med, 16(7), 821–827. doi:10.1038/nm.2167

Bonello, T. T., Perez-Vale, K. Z., Sumigray, K. D., & Peifer, M. (2018). Rap1 acts via multiple mechanisms to position Canoe and adherens junctions and mediate apical-basal polarity establishment. Development, 145(2). doi:10.1242/dev.157941

Bowman, S. K., Neumuller, R. A., Novatchkova, M., Du, Q., & Knoblich, J. A. (2006). The Drosophila NuMA Homolog Mud regulates spindle orientation in asymmetric cell division. Dev Cell, 10(6), 731–742. doi:10.1016/j.devcel.2006.05.005

Buckley, C. D., Tan, J., Anderson, K. L., Hanein, D., Volkmann, N., Weis, W. I., … Dunn, A. R. (2014). Cell adhesion. The minimal cadherin-catenin complex binds to actin filaments under force. Science, 346(6209), 1254211. doi:10.1126/science.1254211

Burridge, K., & Feramisco, J. R. (1982). Alpha-actinin and vinculin from nonmuscle cells: calcium-sensitive interactions with actin. Cold Spring Harb Symp Quant Biol, 46 Pt 2, 587–597.

Byrd, K. M., Lough, K. J., Patel, J. H., Descovich, C. P., Curtis, T. A., & Williams, S. E. (2016). LGN plays distinct roles in oral epithelial stratification, filiform papilla morphogenesis and hair follicle development. Development, 143(15), 2803–2817. doi:10.1242/dev.136010

Cai, Y., Chia, W., & Yang, X. (2001). A family of snail-related zinc finger proteins regulates two distinct and parallel mechanisms that mediate Drosophila neuroblast asymmetric divisions. EMBO J, 20(7), 1704–1714. doi:10.1093/emboj/20.7.1704

Carminati, M., Gallini, S., Pirovano, L., Alfieri, A., Bisi, S., & Mapelli, M. (2016). Concomitant binding of Afadin to LGN and F-actin directs planar spindle orientation. Nat Struct Mol Biol, 23(2), 155–163. doi:10.1038/nsmb.3152

Cetera, M., Leybova, L., Joyce, B., & Devenport, D. (2018). Counter-rotational cell flows drive morphological and cell fate asymmetries in mammalian hair follicles. Nat Cell Biol, 20(5), 541–552. doi:10.1038/s41556-018-0082-7

Cheng, J., Tiyaboonchai, A., Yamashita, Y. M., & Hunt, A. J. (2011). Asymmetric division of cyst stem cells in Drosophila testis is ensured by anaphase spindle repositioning. Development, 138(5), 831–837. doi:10.1242/dev.057901

Choi, H. J., Pokutta, S., Cadwell, G. W., Bobkov, A. A., Bankston, L. A., Liddington, R. C., & Weis, W. I. (2012). alphaE-catenin is an autoinhibited molecule that coactivates vinculin. Proc Natl Acad Sci U S A, 109(22), 8576–8581. doi:10.1073/pnas.1203906109

Choi, W., Acharya, B. R., Peyret, G., Fardin, M. A., Mege, R. M., Ladoux, B., … Peifer, M. (2016). Remodeling the zonula adherens in response to tension and the role of afadin in this response. J Cell Biol, 213(2), 243–260. doi:10.1083/jcb.201506115

Choi, W., Harris, N. J., Sumigray, K. D., & Peifer, M. (2013). Rap1 and Canoe/afadin are essential for establishment of apical-basal polarity in the Drosophila embryo. Mol Biol Cell, 24(7), 945–963. doi:10.1091/mbc.E12-10-0736

Dassule, H. R., Lewis, P., Bei, M., Maas, R., & McMahon, A. P. (2000). Sonic hedgehog regulates growth and morphogenesis of the tooth. Development, 127(22), 4775–4785.

Dor-On, E., Raviv, S., Cohen, Y., Adir, O., Padmanabhan, K., & Luxenburg, C. (2017). T-plastin is essential for basement membrane assembly and epidermal morphogenesis. Sci Signal, 10(481). doi:10.1126/scisignal.aal3154

Du, Q., & Macara, I. G. (2004). Mammalian Pins is a conformational switch that links NuMA to heterotrimeric G proteins. Cell, 119(4), 503–516. doi:10.1016/j.cell.2004.10.028

Gao, L., Yang, Z., Hiremath, C., Zimmerman, S. E., Long, B., Brakeman, P. R., … Marciano, D. K. (2017). Afadin orients cell division to position the tubule lumen in developing renal tubules. Development, 144(19), 3511–3520. doi:10.1242/dev.148908

Geiger, B. (1979).A 130K protein from chicken gizzard: its localization at the termini of microfilament bundles in cultured chicken cells. Cell, 18(1), 193–205.

Geldmacher-Voss, B., Reugels, A. M., Pauls, S., & Campos-Ortega, J. A. (2003). A 90-degree rotation of the mitotic spindle changes the orientation of mitoses of zebrafish neuroepithelial cells. Development, 130(16), 3767–3780.

Gloerich, M., Bianchini, J. M., Siemers, K. A., Cohen, D. J., & Nelson, W. J. (2017). Cell division orientation is coupled to cell-cell adhesion by the E-cadherin/LGN complex. Nat Commun, 8, 13996. doi:10.1038/ncomms13996

Hansen, S. D., Kwiatkowski, A. V., Ouyang, C. Y., Liu, H., Pokutta, S., Watkins, S. C., … Nelson, W. J. (2013). alphaE-catenin actin-binding domain alters actin filament conformation and regulates binding of nucleation and disassembly factors. Mol Biol Cell, 24(23), 3710–3720. doi:10.1091/mbc.E13-07-0388

Hart, K. C., Tan, J., Siemers, K. A., Sim, J. Y., Pruitt, B. L., Nelson, W. J., & Gloerich, M. (2017). E-cadherin and LGN align epithelial cell divisions with tissue tension independently of cell shape. Proc Natl Acad Sci U S A, 114(29), E5845–E5853. doi:10.1073/pnas.1701703114

Haydar, T. F., Ang, E., Jr., & Rakic, P. (2003). Mitotic spindle rotation and mode of cell division in the developing telencephalon. Proc Natl Acad Sci U S A, 100(5), 2890–2895. doi:10.1073/pnas.0437969100

Huang, D. L., Bax, N. A., Buckley, C. D., Weis, W. I., & Dunn, A. R. (2017). Vinculin forms a directionally asymmetric catch bond with F-actin. Science, 357(6352), 703–706. doi:10.1126/science.aan2556

Hyman, A. A., & White, J. G. (1987). Determination of cell division axes in the early embryogenesis of Caenorhabditis elegans. J Cell Biol, 105(5), 2123–2135.

Izumi, Y., Ohta, N., Hisata, K., Raabe, T., & Matsuzaki, F. (2006). Drosophila Pins-binding protein Mud regulates spindle-polarity coupling and centrosome organization. Nat Cell Biol, 8(6), 586–593. doi:10.1038/ncb1409

Kaltschmidt, J. A., Davidson, C. M., Brown, N. H., & Brand, A. H. (2000). Rotation and asymmetry of the mitotic spindle direct asymmetric cell division in the developing central nervous system. Nat Cell Biol, 2(1), 7–12. doi:10.1038/71323

Knoblich, J. A. (2008). Mechanisms of asymmetric stem cell division. Cell, 132(4), 583–597. doi:10.1016/j.cell.2008.02.007

Knoblich, J. A. (2010). Asymmetric cell division: recent developments and their implications for tumour biology. Nat Rev Mol Cell Biol, 11(12), 849–860. doi:10.1038/nrm3010

Komura, H., Ogita, H., Ikeda, W., Mizoguchi, A., Miyoshi, J., & Takai, Y. (2008). Establishment of cell polarity by afadin during the formation of embryoid bodies. Genes Cells, 13(1), 79–90. doi:10.1111/j.1365-2443.2007.01150.x

Kraut, R., Chia, W., Jan, L. Y., Jan, Y. N., & Knoblich, J. A. (1996). Role of inscuteable in orienting asymmetric cell divisions in Drosophila. Nature, 383(6595), 50–55. doi:10.1038/383050a0

Lazaro-Dieguez, F., & Musch, A. (2017). Cell-cell adhesion accounts for the different orientation of columnar and hepatocytic cell divisions. J Cell Biol, 216(11), 3847–3859. doi:10.1083/jcb.201608065

Lechler, T., & Fuchs, E. (2005). Asymmetric cell divisions promote stratification and differentiation of mammalian skin. Nature, 437(7056), 275–280. doi:10.1038/nature03922

Luxenburg, C., Heller, E., Pasolli, H. A., Chai, S., Nikolova, M., Stokes, N., & Fuchs, E. (2015). Wdr1-mediated cell shape dynamics and cortical tension are essential for epidermal planar cell polarity. Nat Cell Biol, 17(5), 592–604. doi:10.1038/ncb3146

Mandai, K., Nakanishi, H., Satoh, A., Obaishi, H., Wada, M., Nishioka, H., … Takai, Y. (1997). Afadin: A novel actin filament-binding protein with one PDZ domain localized at cadherin-based cell-to-cell adherens junction. J Cell Biol, 139(2), 517–528.

Martin-Belmonte, F., & Perez-Moreno, M. (2011). Epithelial cell polarity, stem cells and cancer. Nat Rev Cancer, 12(1), 23–38. doi:10.1038/nrc3169

McKinley, K. L., Stuurman, N., Royer, L. A., Schartner, C., Castillo-Azofeifa, D., Delling, M., … Vale, R. D. (2018). Cellular aspect ratio and cell division mechanics underlie the patterning of cell progeny in diverse mammalian epithelia. Elife, 7. doi:10.7554/eLife.36739

Miroshnikova, Y. A., Le, H. Q., Schneider, D., Thalheim, T., Rubsam, M., Bremicker, N., … Wickstrom, S. A. (2018). Adhesion forces and cortical tension couple cell proliferation and differentiation to drive epidermal stratification. Nat Cell Biol, 20(1), 69–80. doi:10.1038/s41556-017-0005-z

Mora-Bermudez, F., Matsuzaki, F., & Huttner, W. B. (2014). Specific polar subpopulations of astral microtubules control spindle orientation and symmetric neural stem cell division. Elife, 3. doi:10.7554/eLife.02875

Neumuller, R. A., & Knoblich, J. A. (2009). Dividing cellular asymmetry: asymmetric cell division and its implications for stem cells and cancer. Genes Dev, 23(23), 2675–2699. doi:10.1101/gad.1850809

Noethel, B., Ramms, L., Dreissen, G., Hoffmann, M., Springer, R., Rubsam, M., … Hoffmann, B. (2018). Transition of responsive mechanosensitive elements from focal adhesions to adherens junctions on epithelial differentiation. Mol Biol Cell, 29(19), 2317–2325. doi:10.1091/mbc.E17-06-0387

Ouspenskaia, T., Matos, I., Mertz, A. F., Fiore, V. F., & Fuchs, E. (2016). WNT-SHH Antagonism Specifies and Expands Stem Cells prior to Niche Formation. Cell, 164(1-2), 156–169. doi:10.1016/j.cell.2015.11.058

Peng, C. Y., Manning, L., Albertson, R., & Doe, C. Q. (2000). The tumour-suppressor genes lgl and dlg regulate basal protein targeting in Drosophila neuroblasts. Nature, 408(6812), 596–600. doi:10.1038/35046094

Pokutta, S., Drees, F., Takai, Y., Nelson, W. J., & Weis, W. I. (2002). Biochemical and structural definition of the l-afadin- and actin-binding sites of alpha-catenin. J Biol Chem, 277(21), 18868–18874. doi:10.1074/jbc.M201463200

Poulson, N. D., & Lechler, T. (2010). Robust control of mitotic spindle orientation in the developing epidermis. J Cell Biol, 191(5), 915–922. doi:10.1083/jcb.201008001

Rakotomamonjy, J., Brunner, M., Juschke, C., Zang, K., Huang, E. J., Reichardt, L. F., & Chenn, A. (2017). Afadin controls cell polarization and mitotic spindle orientation in developing cortical radial glia. Neural Dev, 12(1), 7. doi:10.1186/s13064-017-0085-2

Ratheesh, A., & Yap, A. S. (2012). A bigger picture: classical cadherins and the dynamic actin cytoskeleton. Nat Rev Mol Cell Biol, 13(10), 673–679. doi:10.1038/nrm3431

Rebollo, E., Roldan, M., & Gonzalez, C. (2009). Spindle alignment is achieved without rotation after the first cell cycle in Drosophila embryonic neuroblasts. Development, 136(20), 3393–3397. doi:10.1242/dev.041822

Rubsam, M., Mertz, A. F., Kubo, A., Marg, S., Jungst, C., Goranci-Buzhala, G., … Niessen, C. M. (2017). E-cadherin integrates mechanotransduction and EGFR signaling to control junctional tissue polarization and tight junction positioning. Nat Commun, 8(1), 1250. doi:10.1038/s41467-017-01170-7

Sawyer, J. K., Choi, W., Jung, K. C., He, L., Harris, N. J., & Peifer, M. (2011). A contractile actomyosin network linked to adherens junctions by Canoe/afadin helps drive convergent extension. Mol Biol Cell, 22(14), 2491–2508. doi:10.1091/mbc.E11-05-0411

Schaefer, M., Shevchenko, A., Shevchenko, A., & Knoblich, J. A. (2000). A protein complex containing Inscuteable and the Galpha-binding protein Pins orients asymmetric cell divisions in Drosophila. Curr Biol, 10(7), 353–362.

Schlegelmilch, K., Mohseni, M., Kirak, O., Pruszak, J., Rodriguez, J. R., Zhou, D., … Camargo, F. D. (2011). Yap1 acts downstream of alpha-catenin to control epidermal proliferation. Cell, 144(5), 782–795. doi:10.1016/j.cell.2011.02.031

Schober, M., Schaefer, M., & Knoblich, J. A. (1999). Bazooka recruits Inscuteable to orient asymmetric cell divisions in Drosophila neuroblasts. Nature, 402(6761), 548–551. doi:10.1038/990135

Siegrist, S. E., & Doe, C. Q. (2005). Microtubule-induced Pins/Galphai cortical polarity in Drosophila neuroblasts. Cell, 123(7), 1323–1335. doi:10.1016/j.cell.2005.09.043

Siller, K. H., Cabernard, C., & Doe, C. Q. (2006). The NuMA-related Mud protein binds Pins and regulates spindle orientation in Drosophila neuroblasts. Nat Cell Biol, 8(6), 594–600. doi:10.1038/ncb1412

Siller, K. H., & Doe, C. Q. (2009). Spindle orientation during asymmetric cell division. Nat Cell Biol, 11(4), 365–374. doi:10.1038/ncb0409-365

Smart, I. H. (1970). Variation in the plane of cell cleavage during the process of stratification in the mouse epidermis. Br J Dermatol, 82(3), 276–282.

Speicher, S., Fischer, A., Knoblich, J., & Carmena, A. (2008). The PDZ protein Canoe regulates the asymmetric division of Drosophila neuroblasts and muscle progenitors. Curr Biol, 18(11), 831–837. doi:10.1016/j.cub.2008.04.072

Vasioukhin, V., Bauer, C., Degenstein, L., Wise, B., & Fuchs, E. (2001). Hyperproliferation and defects in epithelial polarity upon conditional ablation of alpha-catenin in skin. Cell, 104(4), 605–617.

Vasioukhin, V., Bauer, C., Yin, M., & Fuchs, E. (2000). Directed actin polymerization is the driving force for epithelial cell-cell adhesion. Cell, 100(2), 209–219.

Wee, B., Johnston, C. A., Prehoda, K. E., & Doe, C. Q. (2011). Canoe binds RanGTP to promote Pins(TPR)/Mud-mediated spindle orientation. J Cell Biol, 195(3), 369–376. doi:10.1083/jcb.201102130

Weiss, E. E., Kroemker, M., Rudiger, A. H., Jockusch, B. M., & Rudiger, M. (1998). Vinculin is part of the cadherin-catenin junctional complex: complex formation between alpha-catenin and vinculin. J Cell Biol, 141(3), 755–764.

Williams, S. E., Beronja, S., Pasolli, H. A., & Fuchs, E. (2011). Asymmetric cell divisions promote Notch-dependent epidermal differentiation. Nature, 470(7334), 353–358. doi:10.1038/nature09793

Williams, S. E., Ratliff, L. A., Postiglione, M. P., Knoblich, J. A., & Fuchs, E. (2014). Par3-mInsc and Galphai3 cooperate to promote oriented epidermal cell divisions through LGN. Nat Cell Biol, 16(8), 758–769. doi:10.1038/ncb3001

Wodarz, A., Ramrath, A., Kuchinke, U., & Knust, E. (1999). Bazooka provides an apical cue for Inscuteable localization in Drosophila neuroblasts. Nature, 402(6761), 544–547. doi:10.1038/990128

Yamashita, Y. M., Jones, D. L., & Fuller, M. T. (2003). Orientation of asymmetric stem cell division by the APC tumor suppressor and centrosome. Science, 301(5639), 1547–1550. doi:10.1126/science.1087795

Yang, Z., Zimmerman, S., Brakeman, P. R., Beaudoin, G. M., 3rd, Reichardt, L. F., & Marciano, D. K. (2013). De novo lumen formation and elongation in the developing nephron: a central role for afadin in apical polarity. Development, 140(8), 1774–1784. doi:10.1242/dev.087957

Yonemura, S., Wada, Y., Watanabe, T., Nagafuchi, A., & Shibata, M. (2010). alpha-Catenin as a tension transducer that induces adherens junction development. Nat Cell Biol, 12(6), 533–542. doi:10.1038/ncb2055

Zheng, Z., Zhu, H., Wan, Q., Liu, J., Xiao, Z., Siderovski, D. P., & Du, Q. (2010). LGN regulates mitotic spindle orientation during epithelial morphogenesis. J Cell Biol, 189(2), 275–288. doi:10.1083/jcb.200910021

Zigman, M., Cayouette, M., Charalambous, C., Schleiffer, A., Hoeller, O., Dunican, D., … Knoblich, J. A. (2005). Mammalian inscuteable regulates spindle orientation and cell fate in the developing retina. Neuron, 48(4), 539–545. doi:10.1016/j.neuron.2005.09.030

